# Novel allergen discovery through comprehensive *de novo* transcriptomic analyses of 5 shrimp species

**DOI:** 10.1101/2020.06.05.135731

**Authors:** Shaymaviswanathan Karnaneedi, Roger Huerlimann, Elecia B. Johnston, Roni Nugraha, Thimo Ruethers, Aya C. Taki, Sandip D. Kamath, Nicholas M. Wade, Dean R. Jerry, Andreas L. Lopata

**Author notes:** (Lopata AL).

## Abstract

Shellfish allergy affects up to 2% of the world’s population and persists for life in most patients. The diagnosis of a shellfish allergy, in particular shrimp, is however often challenging due to the similarity of allergenic proteins in other invertebrates. Despite the clinical importance, the complete allergen repertoire of allergy-causing shrimps remains unclear. Here we mine the complete transcriptome of five frequently consumed shrimp species to identify and compare allergens with all known allergen sources. The transcriptomes were assembled *de novo* from raw RNA-Seq data of the whiteleg shrimp (*Litopenaeus vannamei*), black tiger shrimp (*Penaeus monodon*), banana shrimp (*Fenneropenaeus merguiensis*), king shrimp (*Melicertus latisulcatus*), and endeavour shrimp (*Metapenaeus endeavouri*). Trinity was used to assemble the transcriptome, and Transrate and BUSCO applied to verify the assembly. Blast search with the two major allergen databases, WHO/IUIS Allergen Nomenclature and AllergenOnline, successfully identified all seven known crustacean allergens. Salmon was utilised to measure their relative abundance, demonstrating sarcoplasmic calcium-binding protein, arginine kinase and myosin light chain as highly abundant allergens. In addition, the analyses revealed up to 40 unreported allergens in different shrimp species, including heat shock protein (HSP), alpha-tubulin, chymotrypsin, cyclophilin, beta-enolase, aldolase A, and glyceraldehyde-3-phosphate dehydrogenase (G3PD). Multiple sequence alignment, conducted in Jalview 2.1 with Clustal Omega, demonstrated high homology with allergens from other invertebrates including mites and cockroaches. This first transcriptomic analyses of allergens in a major food source provides a valuable genomic resource for investigating shellfish allergens, comparing invertebrate allergens and developing improved diagnostics and novel immunotherapeutics for food allergy.

## Introduction

Food allergy affects up to 10% of children and 10% of adults, and the prevalence is projected to be on the rise [1, 2]. Food allergy is caused through ingestion of food that contains allergenic proteins that triggers adverse reactions in sensitised individuals [3, 4]. The term “allergen” refers to a protein capable of inducing sensitisation and subsequent allergic immune responses through immunoglobulin E (IgE)-mediated type 1 hypersensitivity in patients [3–7].

Shellfish allergy is often lifelong, similar to peanut allergy, affects about 2% of the global population and appears to be highly prevalent in the Asia-Pacific region and other countries where seafood consumption is high [8–11]. A recent epidemiology study from Vietnam revealed that the prevalence of shellfish allergy is as high as 4.2% [11], while up to 3% of adults in the USA are sensitised to shellfish [1].

Among shellfish allergic individuals, shrimp allergy seems to be the most prominent crustacean allergy and remains to be difficult to diagnose and manage, for multiple reasons. Shrimp accounts for one of the most prevalent events of food derived anaphylactic reactions after peanuts and tree nuts [12–15].

The management of shrimp allergy is often challenging due to immunological cross-reactivity to molecularly similar allergens [12, 13, 16–22]. The similarity of shrimp allergens to proteins of other shellfish species, including crabs and lobsters, and other invertebrates such as house dust mites (HDM) and cockroaches, can induce unexpected allergic reactions [15–20, 23, 24]. Although this cross-reactivity has been observed in the clinical setting, the underlying molecular basis is not well understood.

Over the past decades, more than 2000 allergens are now well characterised and accessible via several databases, including the World Health Organization & International Union of Immunological Societies (WHO/IUIS) Allergen Nomenclature database (www.allergen.org) having the most stringent inclusion criteria [7, 25], and the highly peer-reviewed AllergenOnline: Home of the FARRP (Food Allergy Research and Resource Program) Allergen Protein database (www.allergenonline.org) [26, 27].

Allergen discovery is traditionally conducted using whole allergen sources and isolation of IgE antibody binding proteins [4, 28–31]. However, this approach has many limitations, including low sensitivity and small patient cohorts that does not allow the detection of all possible allergenic proteins [32].

Here we report the first complete transcriptome analysis of shellfish food allergen source, with focus on shrimp allergens, the most common shellfish allergy. The transcriptomes of five most frequently consumed shrimp species were assembled *de novo* and screened for the presence of similar amino acid (AA) sequences to 2,172 allergens in the WHO/IUIS Allergen Nomenclature and AllergenOnline databases (Figure 1).

**Figure 1:**
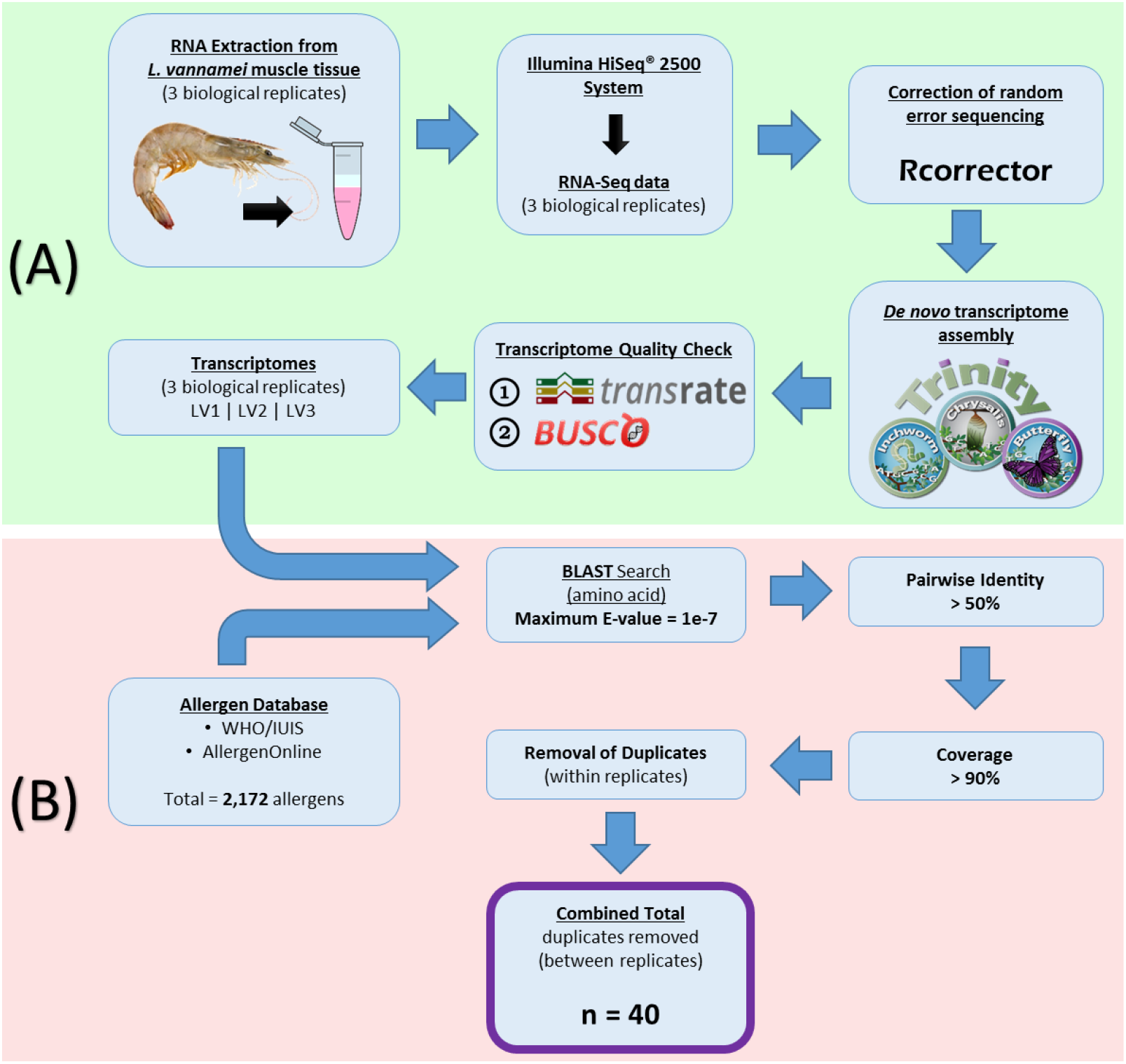
**Schematic representation of (A) *de novo* transcriptome assembly and (B) transcriptomic analysis used in the identification of allergens in shrimps.** The example shown here is for *L. vannamei* species. LV1, LV2, and LV3 represents the 3 biological replicates of *L. vannamei* samples. ‘n’ value refer to the number of allergens identified in *L. vannamei*. A total of 40 allergens were identified.

## Results

### Assessment of 15 assembled transcriptomes

Illumina HiSeq® 2500 (Illumina Australia and New Zealand, VIC, Australia) sequencing produced 125bp paired-end sequencing data with a total number of paired-end reads for each sample of approximately 20 million reads. The *de novo* assembly for 15 samples (three replicates each for five shrimp species) resulted in 28,101 to 42,510 contigs (Table 1). All 15 samples had more than 87% of read pairs that mapped back to the contigs within the assembled transcriptome. TransRate scores (assembly scores) for each of the 15 transcriptomes were approximately 0.4 (Table 1). BUSCO (Benchmarking Universal Single-Copy Orthologs) results, overall, had a complete genes (C) score ranging between 43% – 67%; fragmented genes (F) score ranging between 16% – 26%; and missing genes (M) score between 14% – 32% (Table 1). The transcriptomes of *P. monodon* and *F. merguiensis* had the highest values for complete BUSCO’s (C scores) (Table 1). An observable pattern here is that both these shrimp species also had the highest number of contigs and assembly size.

**Table 1:**
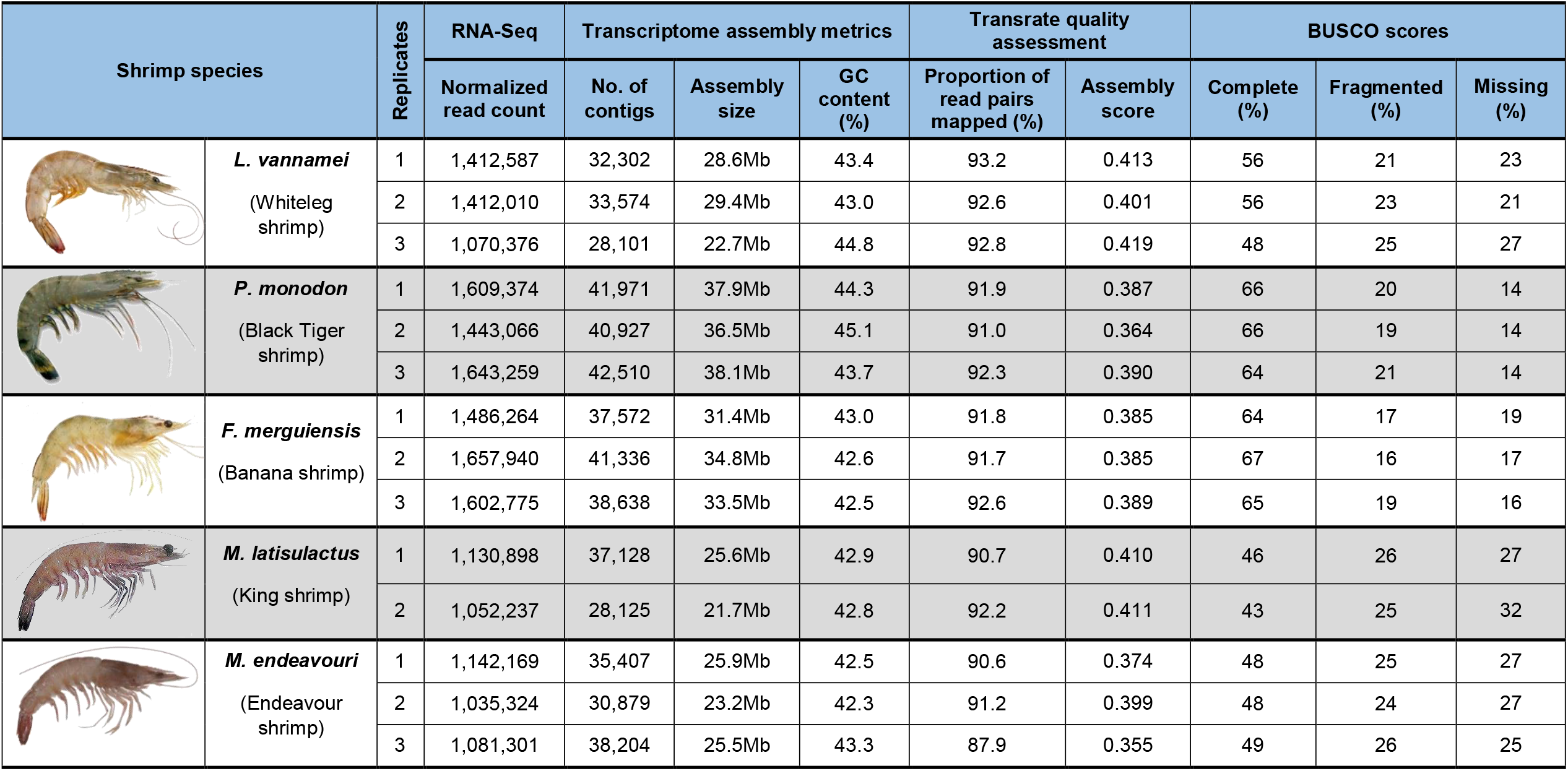
Results of Trinity transcriptome assembly, TransRate, and BUSCO. Shrimp species name (common name) and their 1-3 biological replicates are shown here with their transcriptomes’ number of contigs and assembly size after assembly by Trinity. TransRate score and BUSCO scores (C: complete, F: fragmented, M: missing) of each transcriptome are also shown here.

| Shrimp species |  | Replicates | RNA-Seq | Transcriptome assembly metrics |  |  | Transrate quality assessment |  | BUSCO scores |  |  |
| --- | --- | --- | --- | --- | --- | --- | --- | --- | --- | --- | --- |
|  |  |  | Normalized read count | No. of contigs | Assembly size | GC content (%) | Proportion of read pairs mapped (%) | Assembly score | Complete (%) | Fragmented (%) | Missing (%) |
|  | ***L. vannamei***<br>(Whiteleg shrimp) | 1 | 1,412,587 | 32,302 | 28.6Mb | 43.4 | 93.2 | 0.413 | 56 | 21 | 23 |
|  |  | 2 | 1,412,010 | 33,574 | 29.4Mb | 43.0 | 92.6 | 0.401 | 56 | 23 | 21 |
|  |  | 3 | 1,070,376 | 28,101 | 22.7Mb | 44.8 | 92.8 | 0.419 | 48 | 25 | 27 |
|  | ***P. monodon***<br>(Black Tiger shrimp) | 1 | 1,609,374 | 41,971 | 37.9Mb | 44.3 | 91.9 | 0.387 | 66 | 20 | 14 |
|  |  | 2 | 1,443,066 | 40,927 | 36.5Mb | 45.1 | 91.0 | 0.364 | 66 | 19 | 14 |
|  |  | 3 | 1,643,259 | 42,510 | 38.1Mb | 43.7 | 92.3 | 0.390 | 64 | 21 | 14 |
|  | ***F. merguensis***<br>(Banana shrimp) | 1 | 1,486,264 | 37,572 | 31.4Mb | 43.0 | 91.8 | 0.385 | 64 | 17 | 19 |
|  |  | 2 | 1,657,940 | 41,336 | 34.8Mb | 42.6 | 91.7 | 0.385 | 67 | 16 | 17 |
|  |  | 3 | 1,602,775 | 38,638 | 33.5Mb | 42.5 | 92.6 | 0.389 | 65 | 19 | 16 |
|  | ***M. latisulactus***<br>(King shrimp) | 1 | 1,130,898 | 37,128 | 25.6Mb | 42.9 | 90.7 | 0.410 | 46 | 26 | 27 |
|  |  | 2 | 1,052,237 | 28,125 | 21.7Mb | 42.8 | 92.2 | 0.411 | 43 | 25 | 32 |
|  | ***M. endeavouri***<br>(Endeavour shrimp) | 1 | 1,142,169 | 35,407 | 25.9Mb | 42.5 | 90.6 | 0.374 | 48 | 25 | 27 |
|  |  | 2 | 1,035,324 | 30,879 | 23.2Mb | 42.3 | 91.2 | 0.399 | 48 | 24 | 27 |
|  |  | 3 | 1,081,301 | 38,204 | 25.5Mb | 43.3 | 87.9 | 0.355 | 49 | 26 | 25 |

### Large numbers of allergens identified within the transcriptomes

After duplicate removal, the results yielded 40 unique allergen (AA sequences) identified in whiteleg shrimp (*L. vannamei*), 44 in black tiger shrimp (*P. monodon*), 42 in banana shrimp (*F. merguiensis*), 44 in king shrimp (*M. latisulcatus*), and 50 in endeavour shrimp (*M. endeavouri*) (Figure 2). Approximately two thirds of allergen AA sequences that matched with all five shrimp species’ transcriptomes, belonged to shellfish, mites, and fungi species (Figure 2). The remaining allergen AA sequences belonged to plants, insects, fish and other species.

**Figure 2:**
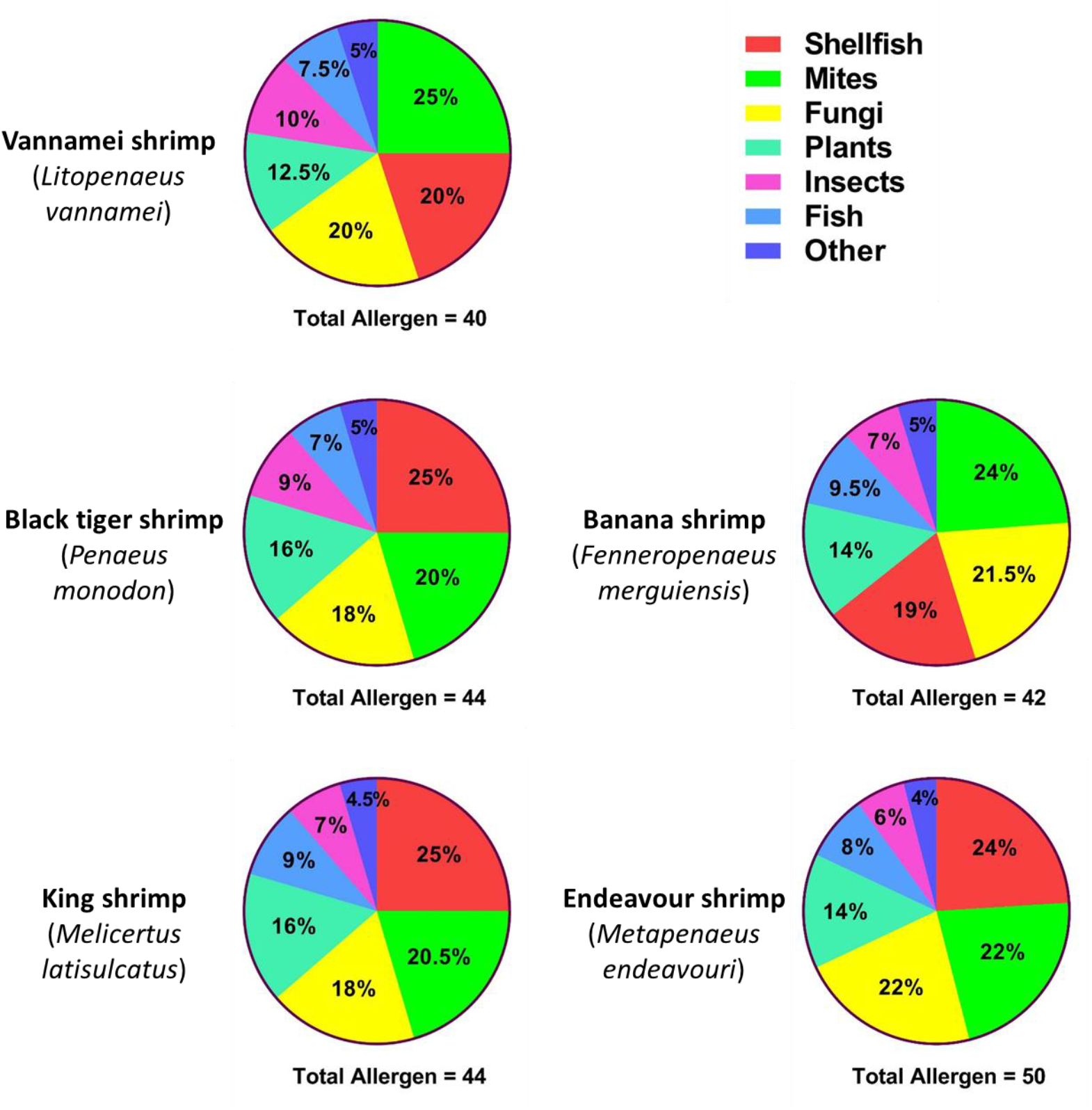
Total allergens identified from the transcriptomic analysis in each five shrimp species, distributed based on the matched allergen’s source. The distribution amongst different groups of allergen sources are shown in percentages and arranged in a descending order.

### Known crustacean allergens identified

Contigs that matched with the major shrimp allergen tropomyosin (TM) were identified in all five species, with some species having more than one contig representing this allergen (Figure 3.A). *L. vannamei* shrimp’s TM_Contig_1 has a 100% AA sequence identity with the previously recorded and IUIS registered Lit v 1 (ACB38288). This is a similar finding to *P. monodon’*s TM_Contig_1, which has a 100% sequence identity with Pen m 1 (AAX37288). Both TM_Contig_1’s also match with a 100% similarity with each other (Figure 3.A). Overall, TM_Contig_1 of all five species showed a high sequence similarity (pairwise identity of 99-100%) with Lit v 1 and Pen m 1, (Figure 3.A). However, TM_Contig_2 of *P. monodon* and *M. endeavouri* only showed a pairwise identity (PI) of 91% and 82%, respectively, with both Lit v 1 and Pen m 1 (Figure 3.A). The inclusion of HDM and cockroach tropomyosin allergens, Der p 10 (AAB69424), Bla g 7 (AAF72534), and Per a 7 (CAB38086), in the analyses of tropomyosin AA sequences revealed to have more than 70% PI with shrimp TM (Figure 3.A). Molecular phylogenetic tree analyses of previously published AA sequences of TM revealed that the crustacean TM’s are not only very similar to each other, but also to insect and mite TM’s. In comparison, molluscs, which are also grouped as “shellfish” with crustaceans, were found to be only distantly related in terms of TM AA sequence (Figure 3.B).

**Figure 3:**
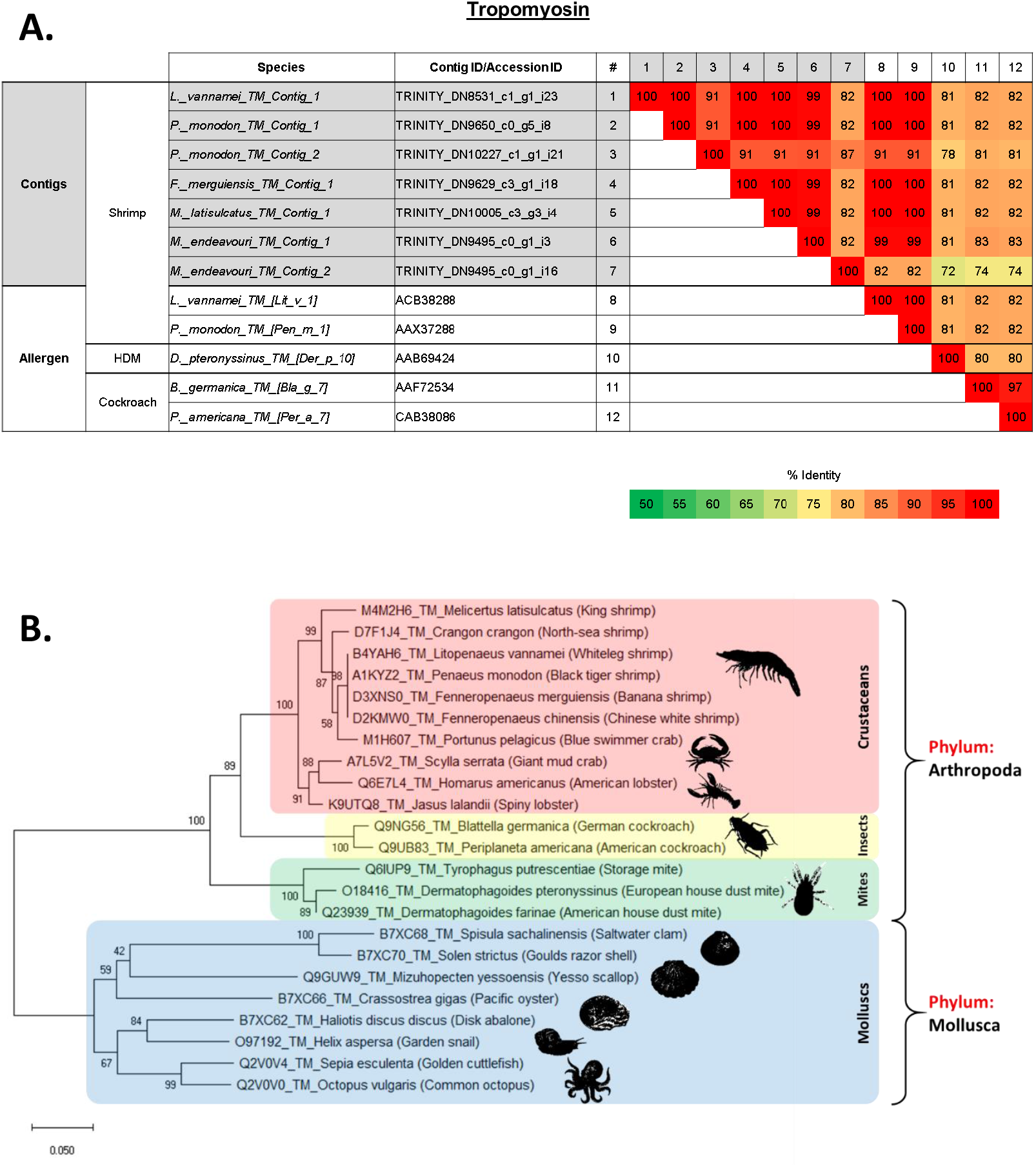
**A. Comparison of amino acid sequence identities of (1-7) contigs from five shrimp species that matched with tropomyosin (TM) allergen, (8-9) known shrimp TM allergen, and (10-12) house dust mite and cockroach TM allergen.** The sequence identities were calculated using multiple sequence alignment in Clustal Omega (EMBL-EBI). **B. Molecular phylogenetic tree based on published amino acid sequences of Tropomyosin (TM) from edible crustacean and mollusc species; and allergy causing mite and insect species.** The branches consist of UniProt ID/Genbank Accesion ID, species name, and followed by common name in brackets. The numbers next to the branches indicate the bootstrap test percentage of 10,000 replicate trees.

One contig in each of four shrimp species was identified as the arginine kinase (AK) allergen, while *M. endeavouri* had two contigs. In contrast to TM, all contigs were highly similar to each other and to the published AK allergens in *L. vannamei*, Lit v 2 (ABI98020), and *P. monodon*, Pen m 2 (AAO15713), with more than 95% PI (Figure 4.A). They were all also found to be more similar to the published cockroach AK allergens in *B. germanica*, Bla g 9 (ACM24358), and *P. americana*, Per a 9 (AAT77152), (83-84% identity) than the published HDM AK allergens in *D. pteronyssinus*, Der p 20 (ACD50950), and *D. farinae*, Der f 20 (AIO08850) (78-79% identity) (Figure 4.A). Similar to TM, published AA sequences of crustacean AK are more closely related to each other; and insects and mites as opposed to molluscs (Figure 4.B).

**Figure 4:**
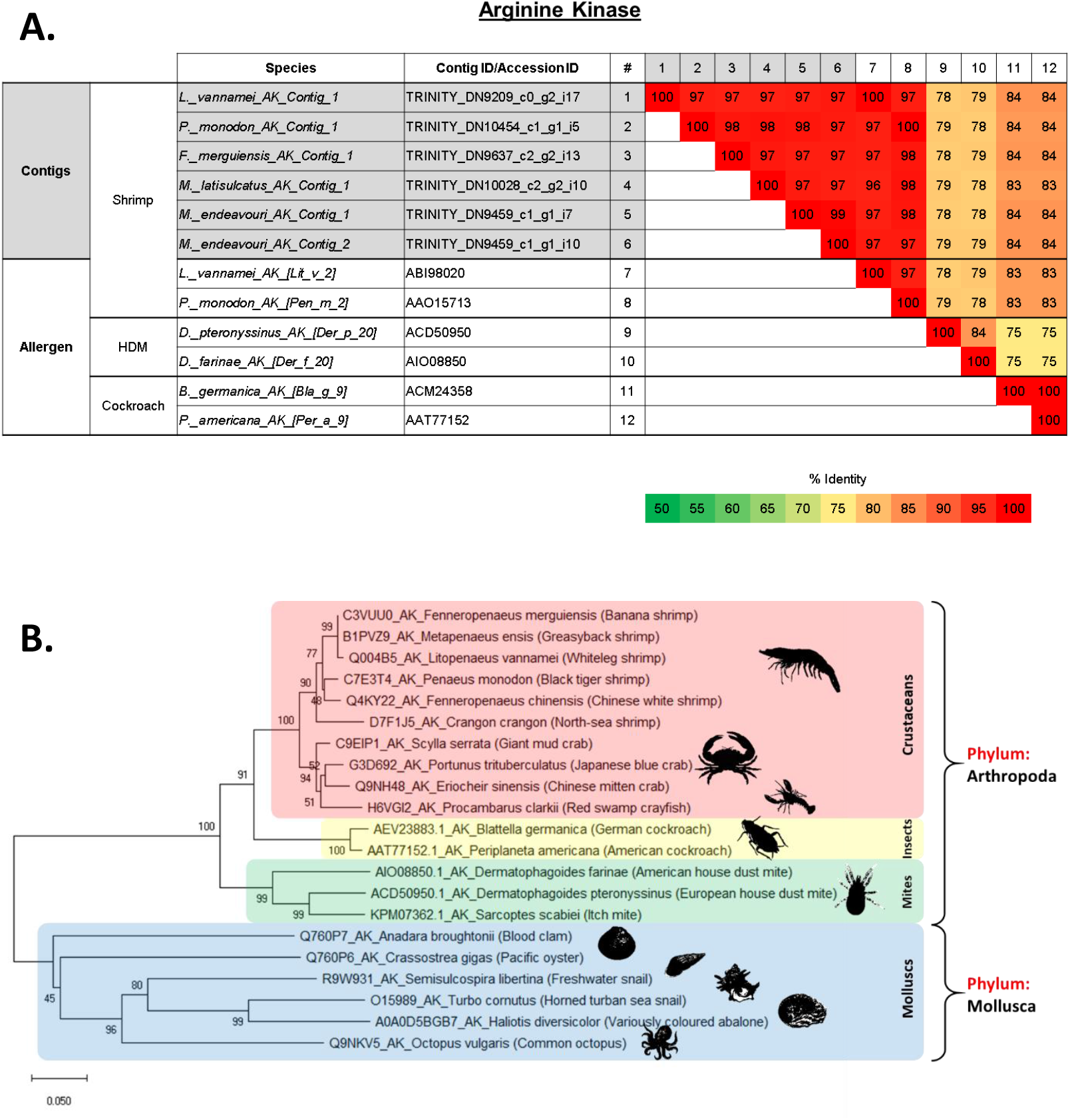
**A. Comparison of amino acid sequence identities of (1-6) contigs from five shrimp species that matched with arginine kinase (AK) allergen (7-8) known shrimp AK allergen and (9-12) house dust mite and cockroach AK allergen.** The sequence identities were calculated using multiple sequence alignment in Clustal Omega (EMBL-EBI). **B. Molecular phylogenetic tree based on published amino acid sequences of Arginine kinase (AK) from edible crustacean and mollusc species; and allergy causing mite and insect species.** The branches consist of UniProt ID/Genbank Accesion ID, species name, and followed by common name in brackets. The numbers next to the branches indicate the bootstrap test percentage of 10,000 replicate trees.

Only one contig each from all five shrimp species analysed matched myosin light chain (MLC), and demonstrated almost identical AA sequences. Interestingly, they were not at all similar to the published MLC allergens in *L. vannamei*, Lit v 3 (ACC76803), or *P. monodon*, Pen m 3 (ADV17342), with only 16-17% PI (Figure 5.A). Instead, they were found to be more similar to the *C. crangon* (North-sea shrimp) MLC allergen, Cra c 5 (ACR43477) with 86-87% PI (Figure 5.A). The contigs were also identified to be more closely related to the american HDM, *D. farinae*, MLC allergen, Der f 26, (51-54% identity) than the German cockroach, *B. germanica*, MLC allergen, Bla g 8 (18-19% identity) (Figure 5.A). Molecular phylogenetic tree analyses on the distance of MLC among edible crustaceans, molluscs and allergy causing mites confirmed that not all crustacean MLC are closely related to each other. For example, mud crab (*S. paramamosain*) is more closely related to molluscs’ MLC than shrimps and crayfish (Figure 5.B). Black tiger shrimp (*P. monodon*) and whiteleg shrimp (*L. vannamei*) contain MLC that are distantly related to kuruma shrimp (*M. japonicus*) and north-sea shrimp (*C. crangon*), but closely related to MLC from German cockroach (*B. germanica*) (Figure 5.B).

**Figure 5:**
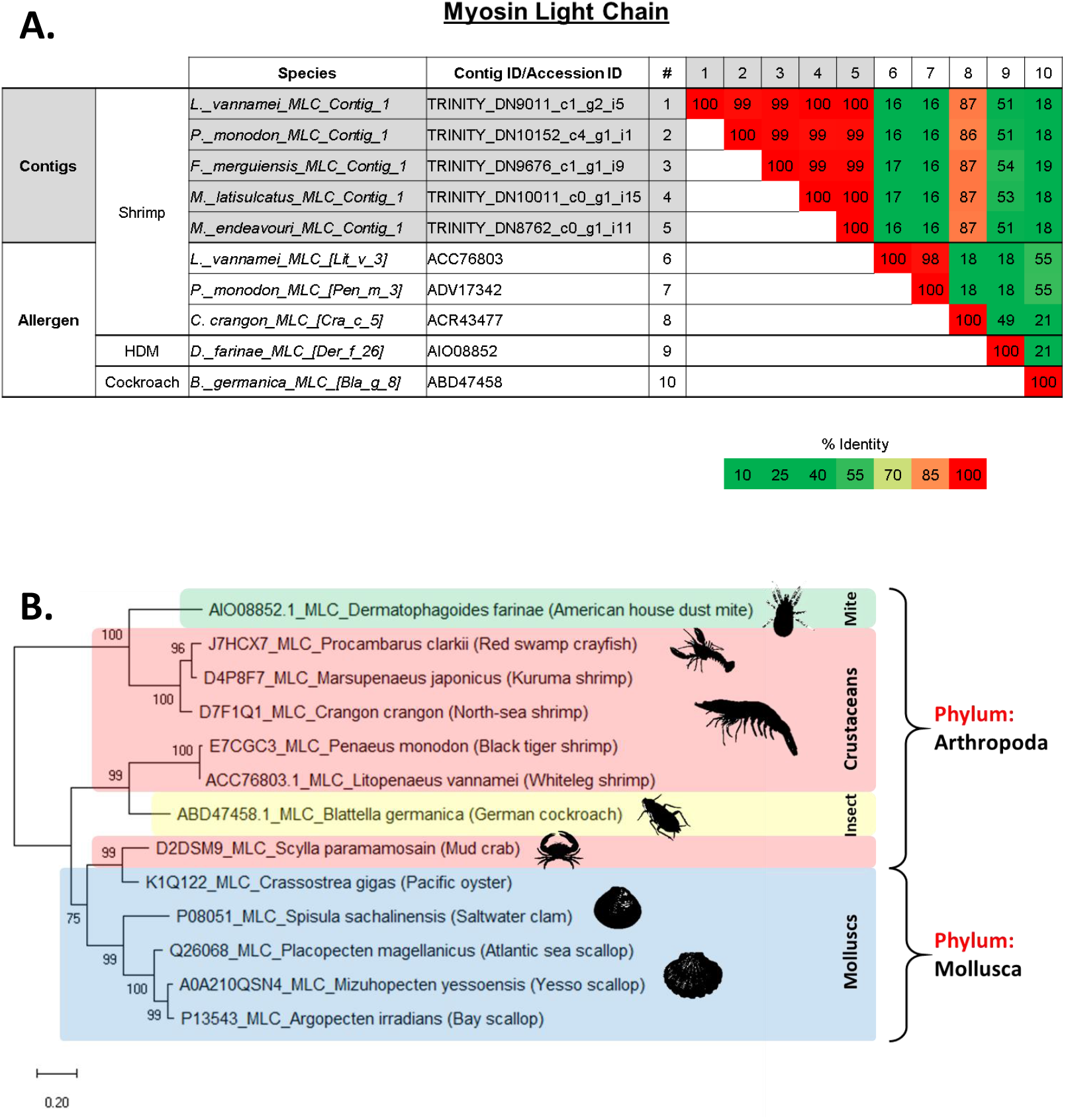
**A. Comparison of amino acid sequence identities of (1-5) contigs from five shrimp species that matched with myosin light chain (MLC) allergen, (6-8) known shrimp MLC allergen and (9-10) house dust mite and cockroach MLC allergen.** The sequence identities were calculated using multiple sequence alignment in Clustal Omega (EMBL-EBI). **B. Molecular phylogenetic tree based on published amino acid sequences of Myosin light chain (MLC) from edible crustacean and mollusc species; and allergy causing mite and insect species.** The branches consist of UniProt ID/Genbank Accesion ID, species name, and followed by common name in brackets. The numbers next to the branches indicate the bootstrap test percentage of 10,000 replicate trees.

Four of the shrimp species had two contigs matching sarcoplasmic calcium-binding protein (SCBP), while *M. endeavouri*, only had one. SCBP_Contig_1 from all five shrimp species were highly similar to each other and also with the published SCBP allergen in *L. vannamei*, Lit v 4 (ACM89179) and *P. monodon*, Pen m 4 (ADV17343) with PI close to 100%, but only over 80% with the published SCBP allergen in *C. crangon*, Cra c 4 (ACR43475) (Figure 6.A). In contrast, SCBP_Contig_2 from the four species, except *M. endeavouri*, were only 82-84% identical to Lit v 4, Pen m 4 and Cra c 4, with the latter having a slightly higher match than the two former (Figure 6.A). Unlike MLC, but similar to TM and AK, the published AA sequences of SCBP in a phylogenetic tree analyses portrayed that all SCBP from edible crustaceans and molluscs are very closely related to other species within the same phylum, but distantly related between the phyla (Figure 6.B).

**Figure 6:**
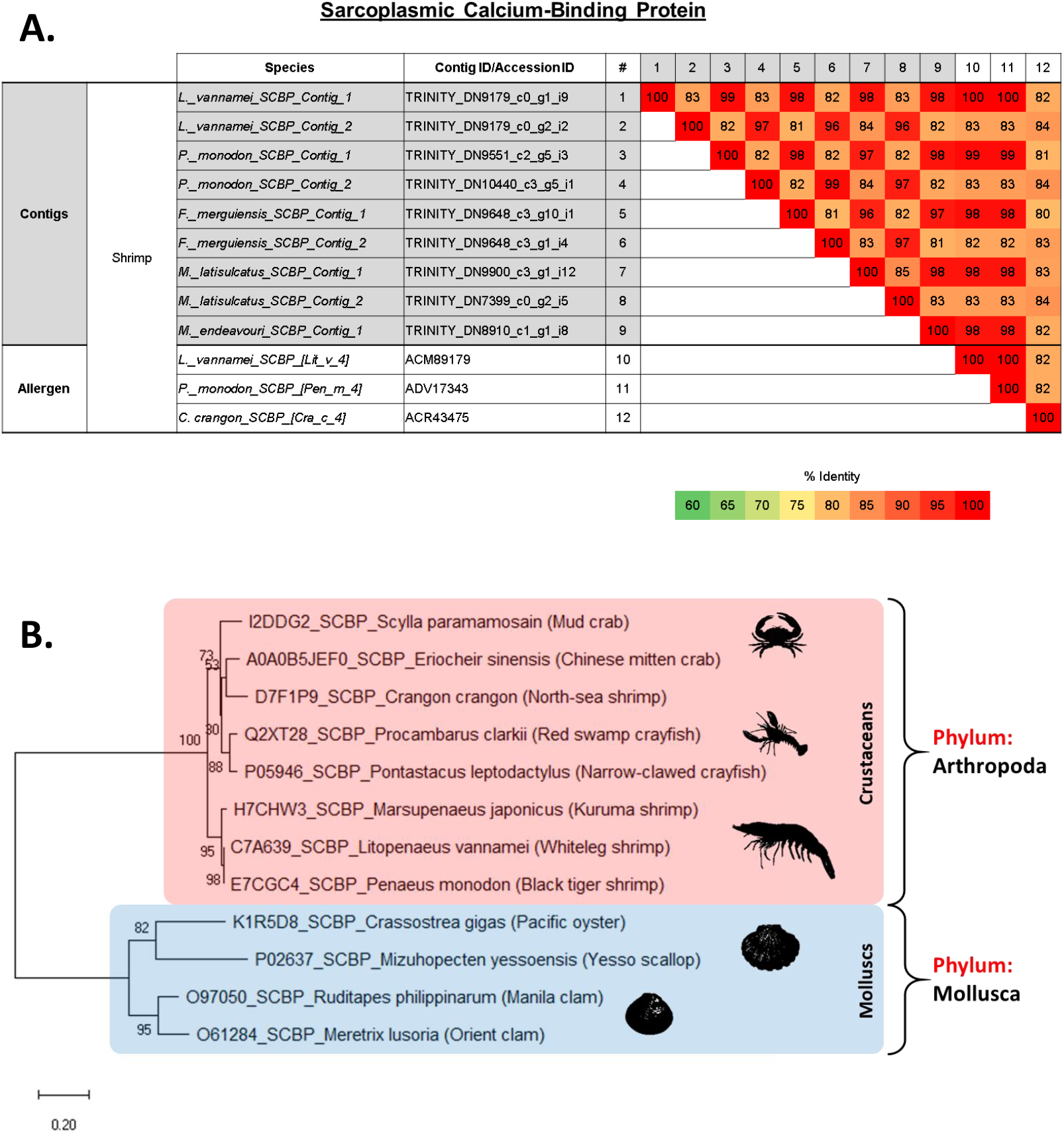
**A. Comparison of amino acid sequence identities of (1-9) contigs from five shrimp species that matched with sarcoplasmic calcium-binding protein (SCBP) allergen and (10-12) known shrimp SCBP allergen.** The sequence identities were calculated using multiple sequence alignment in Clustal Omega (EMBL-EBI). **B. Molecular phylogenetic tree based on published amino acid sequences of Sarcoplasmic calcium-binding protein (SCBP) from edible crustacean and mollusc species; and allergy causing mite and insect species.** The branches consist of UniProt ID/Genbank Accesion ID, species name, and followed by common name in brackets. The numbers next to the branches indicate the bootstrap test percentage of 10,000 replicate trees.

Seven contigs matched with Troponin C (TNC) in all five shrimp species, with *M. latisulcatus* and *M. endeavouri* having two contigs each whilst the other three shrimp species having only one each. All seven contigs were moderate to highly similar to each other and also with the published TNC allergens in *P. monodon*, Pen m 6 (ADV17344) and *C. crangon*, Cra c 6 (ACR43478), with PI ranging between 81-100% (Supplementary Figure 1). The PI of shrimp TNC with cockroach and storage mite TNC allergens ranged between 57-65% (Supplementary Figure 1). Meanwhile, only one contig from each shrimp species matched with Troponin I (TNI) allergen, and they were all highly identical to each other (PI: 87-99%), but were only moderately identical to the published TNI allergen in the narrow-clawed crayfish *P. leptodactylus*, Pon l 7 (P05547) (PI: 78-88%) (Supplementary Figure 2). Similarly, only one contig matched with Triosephosphate isomerase (TIM) allergen in each shrimp species and were all highly identical to each other and also with the published TIM allergen in *C. crangon*, Cra c 8 (ACR43476), (PI: 87-99%) (Supplementary Figure 3). However, they had lower PI to American HDM TIM allergens, Der f 25.01 (AGC56216) and Der f 25.02 (AIO08860), with PI values ranging between 66-69% (Supplementary Figure 3).

### Abundance of known crustacean allergens varies between shrimp species

The average expression or mean abundance, measured in transcripts-per-million (TPM), of TM across all five species ranges from 10,000 – 15,000 TPM (Figure 7.A). Comparing the difference in abundance between the two tropomyosin contigs within the same species (*P. monodon* and *M. endeavouri*), TM_Contig_2 of *P. monodon* was found to be significantly lower than its counterpart, TM_Contig_1 (Figure 7.A). Meanwhile, there was no significant difference between TM_Contig_1 and TM_Contig_2 of *M. endeavouri* (Figure 7.A). With AK, the mean abundance was approximately 40,000 – 80,000 TPM in all five species (Figure 7.B). Comparing the abundance of the two AK contigs in *M. endeavouri*, AK_Contig_1 was significantly lower than AK_Contig_2 (Figure 7.B). The mean abundance of MLC was approximately 30,000 – 50,000 TPM in all species (Figure 7.C). Meanwhile, for SCBP, the mean abundance was between 40,000 and 90,000 TPM in all species (Figure 7.D). Interestingly, SCBP_Contig_1 of *L. vannamei, P. monodon, and F. merguiensis* were all significantly higher than their respective SCBP_Contig_2 (Figure 7.D). The same pattern could be visually observed on *M. latisulcatus* too but unfortunately, due to the presence of only two replicates instead of three (refer to: *Removal of inconclusive dataset* in Materials and methods section), the significance of this difference could not be statistically confirmed by T-test (GraphPad Prism (v7.03)). Similarly, one could not predict the significance of differences in the two TNC contigs of *M. latisulcatus*. However, for *M. endeavouri*, TNC_Contig_1 was found to be significantly higher than its TNC_Contig_2 (Figure 7.E). Overall, the mean abundance value for TNC was around 4,000 – 10,000 TPM for all five shrimp species (Figure 7.E). As for TNI and TIM, the mean abundance values for all five shrimp species were approximately 16,000 – 20,000 TPM (Figure 7.F) and 2,000 – 6,000 TPM respectively (Figure 7.G).

**Figure 7:**
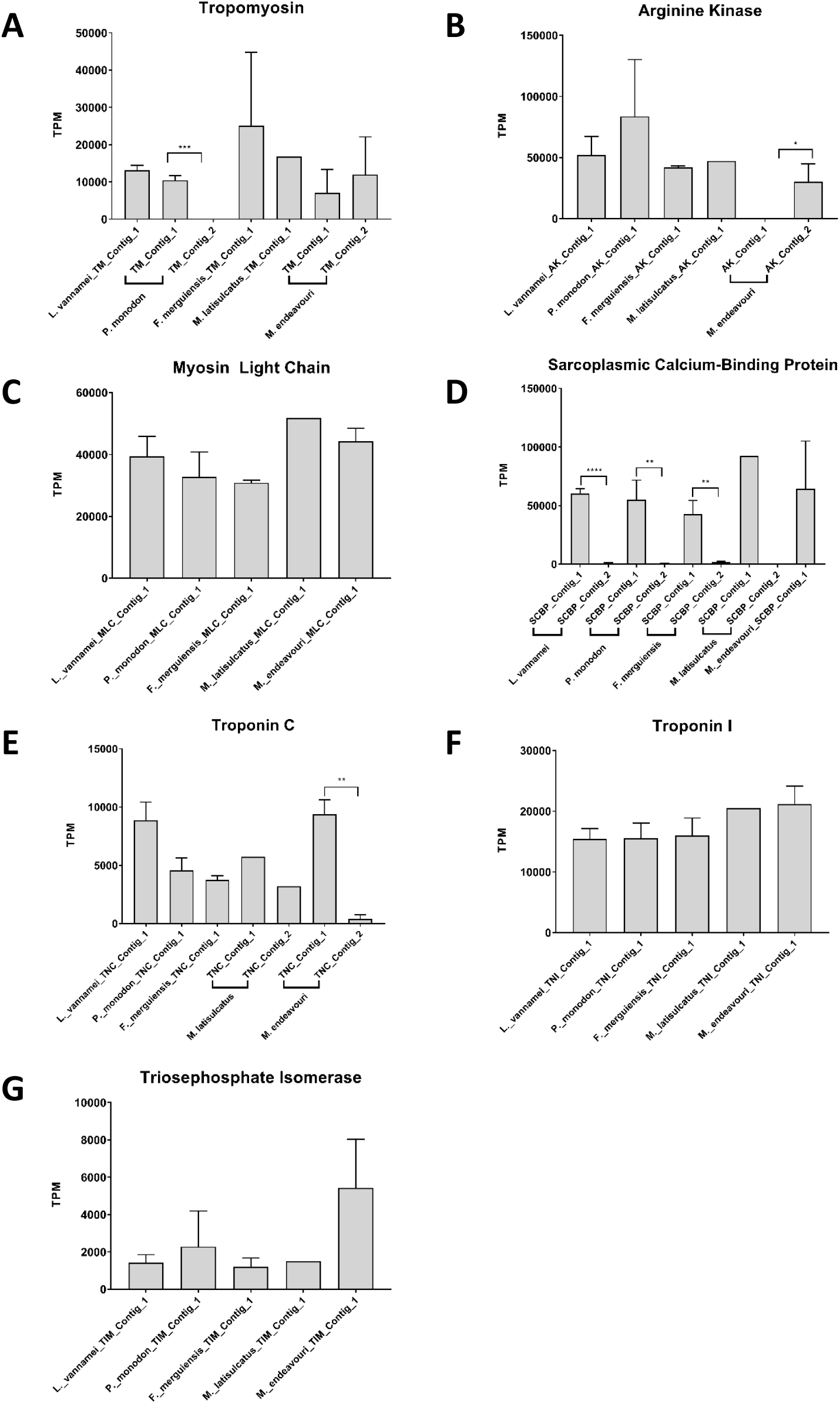
Abundance estimation values in transcript-per-million (TPM) for contigs in the 5 analysed shrimp species that matched with shrimp allergens. **A**: tropomyosin, **B**: arginine kinase, **C**: myosin light chain, **D**: sarcoplasmic calcium-binding protein, **E**: troponin C, **F**: troponin I, and **G**: triosephophate isomerase. *T-test* was employed to measure the significance of difference between two contigs from the same species, if present (*: P ≤ 0.05, **: P ≤ 0.01, ***: P ≤ 0.001, ****: P ≤ 0.0001)

We then examined the difference in abundance of each allergen within individual shrimp species. We only took into account the contig with the highest PI value when there were more than one contig for that allergen. In all species, the top three most highly expressed allergens were SCBP, AK, and MLC (Figure 8). In fact, SCBP was the most highly expressed allergen in all species except *P. monodon*, where AK was higher (Figure 8.B). In descending order of abundance, these three allergens are followed by TNI, TM, TNC, and TIM (Figure 8). However, in *F. merguiensis*, TM was higher than TNI, TNC and TIM (Figure 8.C). In addition, only in *F. merguiensis*, TM’s abundance was not significantly different from all the three highly abundant allergens, namely, SCBP, AK and MLC (Figure 8.C).

**Figure 8:**
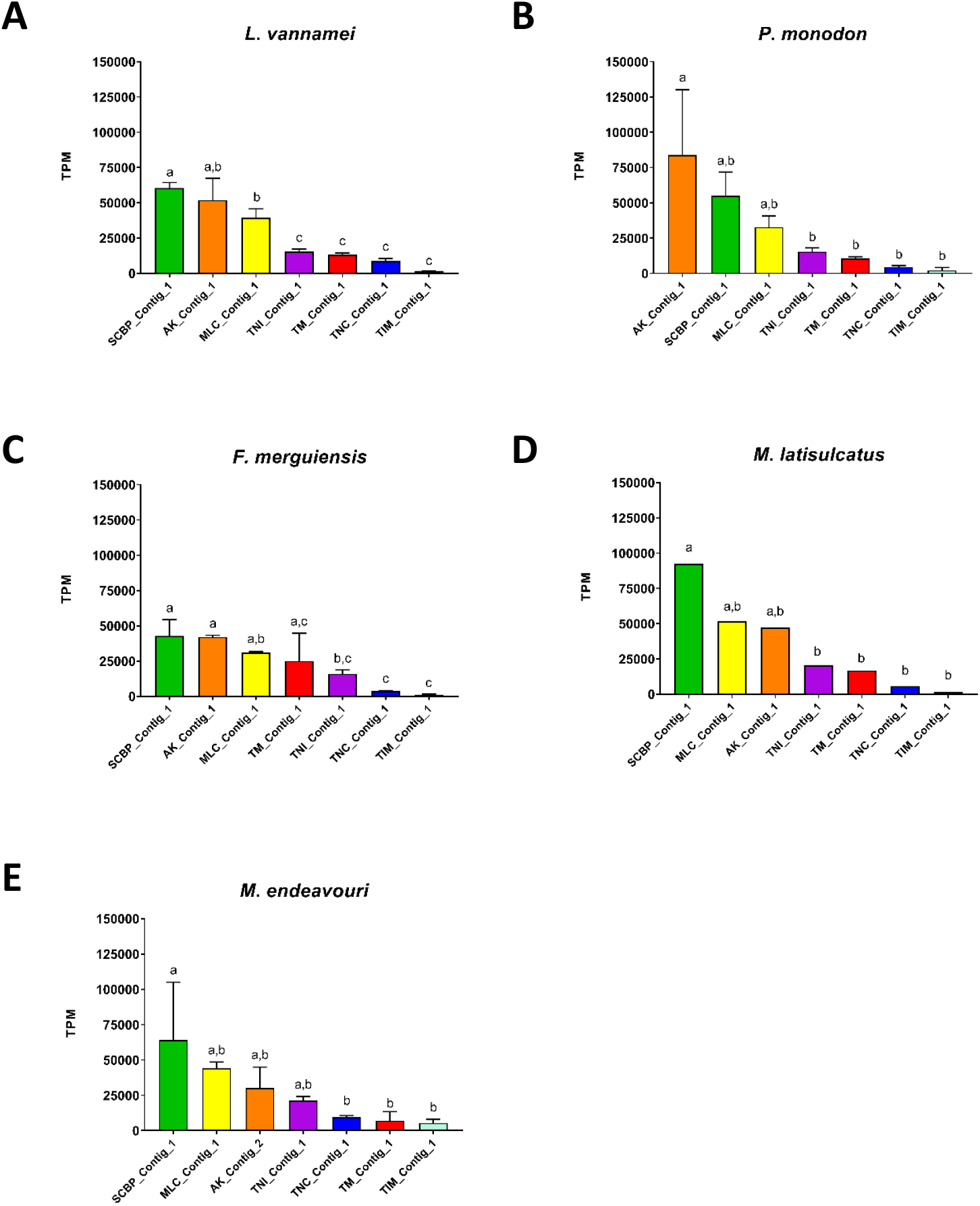
Abundance estimation values in transcript-per-million (TPM) for contigs that matched with shrimp allergens in the 5 analysed shrimp species. **A**: *L. vannamei*, **B**: *P. monodon*, **C**: *F. merguiensis*, **D**: *M. latisulcatus*, **E**: *M. endeavouri*. ANOVA test was employed to measure the significance of difference between the seven shrimp allergens. Only one contig with the highest Pairwise Identity with known shrimp allergens value was included where there was more than one contig for one allergen in each species. The contigs are arranged in descending order of on their abundance. Allergen abundance values with the same letter are not significantly different to each other.

### Evolutionary relationship of shellfish allergens TM, AK, MLC, and SCBP

The evolutionary distance of shrimp TM, AK, MLC, and SCBP were analysed among other edible crustacean and mollusc species; and allergy causing mite and insect species. The generated molecular phylogenies of all four shrimp proteins showed close affinities to homologues of other crustaceans such as crab, lobster and crayfish. However, homologues of the other class of “shellfish”, molluscs, have a distant relationship to shrimps. Molecular phylogenetic analyses of TM and AK revealed that allergy inducing mite and insect homologues are closer to shrimp TM and AK than molluscs. This observation is supported by a recent study by Nugraha et al. where IgE antibody binding epitopes demonstrated shared protein regions of clinical importance [33]. MLC of German cockroach is found to have a closer evolutionary relationship to the black tiger shrimp and whiteleg shrimp, whilst the MLC belonging to American house dust mite is closely related to MLC of a different subset of crustaceans, including, the north-sea shrimp, kuruma shrimp, and red swamp crayfish. Another interesting finding is that the crustacean MLC of mud crab have a closer evolutionary distance with homologues from the mollusca phylum, especially the pacific oyster, but not to those of the other crustaceans. Molecular phylogenetic analysis of SCBP shows a demarcated distance between the crustacean SCBP and mollusc SCBP. No insect or mite SCBP was included in this analyses as there were no AA sequence data available for insect or mite SCBP on NCBI Genbank or UniProt databases.

### Discovery of unreported shrimp allergens

Apart from the previously established shellfish allergens that were confirmed in the five shrimps’ transcriptomes, some of the allergens of non-shellfish species are identified to be standout candidates to be unreported allergens in shrimps due to their high % PI values (Table 2). These allergens are heat shock protein (HSP), alpha-tubulin, chymotrypsin, beta-enolase, glyceraldehyde-3-phosphate dehydrogenase (G3PD), cyclophilin and aldolase A (Table 2). The HSP70 (Tyr p 28, AOD75395) from the storage mite *T. putrescentiae* that matched with the shrimp transcriptomes has the highest PI values with all 5 shrimp species (>82%) (Table 2). Other allergen AA sequences that matched with a PI of more than 70% to all 5 shrimp species’ transcriptomes are alpha-tubulin (Der f 33, AIO08861) and chymotrypsin (Der f 6, AAP35065) of the american HDM *D. farinae*, beta-enolase (Sal s 2, ACH70932) of the atlantic salmon *S. salar*, and glyceraldehyde-3-phosphate dehydrogenase or G3PD (Tri a 34, CAZ76054) of wheat *T. aestivum* (Table 2). The allergen cyclophilin (Asp f 27, CAI78448) of the common mould *A. fumigatus*, only matched with a PI of more than 70% with the banana and king shrimps’ transcriptomes. Meanwhile, the allergen Aldolase A (Thu a 3, CAX62602) of the yellowfin tuna *T. albacares* matched with a PI of more than 70% only with the banana and endeavour shrimps’ transcriptomes (Table 2).

**Table 2:** List of unreported allergens identified that have a minimum of 70% pairwise identity value in at least one species. List includes protein name, the common and scientific name of the allergen source, along with the allergen sequence’s IUIS nomenclature. % Pairwise identity and E-values. Proteins with a % Pairwise identity of 70% or higher (highly likely to be allergenic) are highlighted in red.

| Allergens |  |  |  | LV<br>Whiteleg shrimp<br>(E Value) | PM<br>Black tiger shrimp<br>(E Value) | FM<br>Banana shrimp<br>(E Value) | ML<br>King shrimp<br>(E Value) | ME<br>Endeavour shrimp<br>(E Value) |
| --- | --- | --- | --- | --- | --- | --- | --- | --- |
| Protein name | Source name |  | IUIS<br>nomen-<br>clature |  |  |  |  |  |
|  | Common | Scientific |  |  |  |  |  |  |
| Heat shock-like protein | Storage mite | *Tyrophagus putrescentiae* | Tyr p 28 | 85.1%<br>(0) | 82.7%<br>(0) | 83.3%<br>(0) | 82.7%<br>(0) | 84.3%<br>(0) |
| Alpha-tubulin | American house dust mite | *Dermatophagoides farinae* | Der f 33 | 81.8%<br>(0) | 81.7%<br>(0) | 81.6%<br>(0) | 81.6%<br>(0) | 81.6%<br>(0) |
| Chymotrypsin | American house dust mite | *Dermatophagoides farinae* | Der f 6 | 78.7%<br>(4.3E-94) | 78.7%<br>(2.13E-94) | 79.3%<br>(1.45E-94) | 79.9%<br>(3.71E-97) | 80.5%<br>(3.93E-95) |
| Enolase 3-2 | Atlantic salmon | *Salmo salar* | Sal s 2 | 74.8%<br>(0) | 74.6%<br>(0) | 74.1%<br>(0) | 74.6%<br>(0) | 74.5%<br>(0) |
| Glyceral-dehyde-3-phosphate de-hydrogenase | Wheat | *Triticum aestivum* | Tri a 34 | 72.3%<br>(1.25E-168) | 72.0%<br>(2.87E-172) | 71.7%<br>(1.86E-170) | 72.0%<br>(1.21E-174) | 72.3%<br>(4.91E-175) |
| Cyclophilin | Common mould | *Aspergillus fumigatus* | Asp f 27 | 61.9%<br>(8.45E-65) | 62.5%<br>(1.87E-67) | 70.3%<br>(2.63E-75) | 70.7%<br>(4.38E-75) | 69.3%<br>(5.53E-76) |
| Aldolase A | Yellowfin tuna | *Thunnus albacares* | Thu a 3 | 66.0%<br>(2.29E-164) | 64.9%<br>(2.25E-164) | 70.1%<br>(4.09E-169) | 69.6%<br>(1.47E-166) | 70.1%<br>(3.65E-167) |

## Discussion

Allergen discovery using traditional protein isolation and immunological assay methods, have identified and characterised seven shellfish allergens including the major allergen TM, in addition to AK, MLC, SCBP, TNC, TNI, and TIM. All seven allergens have been identified in various shrimp species except TNI. The increased reporting of allergic cross-reactivity in shrimp-allergic patients to non-shrimp sources demands a full analysis of allergenic proteins. This study utilised an advanced transcriptomic approach to discover the whole repertoire of shrimp allergens, both reported and unreported. Using this comprehensive approach, combining the generation of transcriptomes of five shrimp species and BLAST searching the transcriptomes with all known allergen AA sequences, we identified up to 50 allergens. The majority of identified allergens (45%) belong to the group of shellfish and mite allergens. This is not surprising as the shellfish group consists of crustacean (shrimp) and molluscs, which are often combined when analysing related allergens [15, 33, 34].

In line with existing studies, we confirmed the presence of TM in all five investigated shrimp species, however, the AA sequence was not always similar and the abundance varied significantly. The major allergen TM, a rod-shaped muscle protein (33 – 39 kDa), demonstrates 100% PI between the whiteleg and black tiger shrimp, as recently reported by Ruethers et al. [15], validating the *in silico* approach used in this study. Furthermore, we demonstrated for the first time that TM from banana and king shrimp also exhibited 100% PI to whiteleg and black tiger shrimp’s TM. In contrast, known TM allergen from the king shrimp (Mel l 1; AGF86397) shares only 95% of AA identity with the other shrimp species, which was previously demonstrated to result in different allergenicity in patients [35]. This is an important finding which needs to be followed up in clinical studies. However, the identified TM in all five shrimps are very similar and therefore termed isoallergens. The IUIS Allergen Nomenclature identifies an isoallergen to be two proteins with the same biological function with more than 67% AA sequence identity and similar molecular size [36]. The high AA sequence identity (PI: >70%) of the house dust mite (HDM) and cockroach TM allergens (Der p 10, Bla g 7 and Per a 7) with all the analysed shrimp TM’s indicate a likelihood of all these invertebrate allergens of being cross-reactive. As previously established, an AA sequence identity of more than 70% would demonstrate a highly-likely possibility of cross-reactive IgE antibody binding to these allergens [32, 37]. Clinical studies have previously demonstrated a phenomenon named ‘HDM-cockroach-shrimp’ cross-reactivity [23, 38–42], and we provide here definitive molecular data on the AA sequence similarity of a major shrimp allergen with other invertebrate species.

AK, an important enzymatic protein which regulates the cellular ATP levels of invertebrates, is a heat labile protein (38 – 45 kDa) and highly concentrated in muscle tissue [43]. All five analysed shrimp species demonstrate very high AA sequence similarity with each other, indicating that shrimp allergic patients reacting to AK would most likely react to all five species. AK is considered a major allergen amongst insects and mites and potentially a pan-allergen implicated in cross-reactivity between invertebrate species [44–46]. All five shrimp AK identified in this study are highly-likely allergens with high AA sequence identity (>70%) to AKs from mites (Der p 20; Der f 20) and cockroaches (Bla g 9; Per a 9). Furthermore, the two almost similar AK contigs (PI: 99%) found in endeavour shrimp indicate that they are potential variants instead of isoforms [36], however, the significantly low abundance of AK_Contig_1 than AK_Contig_2 of endeavour shrimp reduces the likelihood of AK_Contig_1 having a role as an allergen.

MLC is part of a large macromolecular complex in muscle tissue consisting of two heavy and four light chains. There are two crustacean MLC allergens, the essential MLC1 (~18kDa) and the regulatory MLC2 (~20kDa) [47, 48]. MLC allergens have been identified in the whiteleg shrimp (Lit v 3) [49] and black tiger shrimp (Pen m 3) [28] – both MLC2 – but not in other shrimp species investigated in this study. While the AA sequences of MLC1 and MLC2 are very different from each other [50], the MLC contigs found in all 5 analysed shrimps are most likely MLC1. Furthermore, this study also suggests that HDM MLC (Der f 26) and cockroach MLC (Bla g 8) are most likely MLC1 and MLC2, respectively, explaining the close molecular phylogenetic relationship to crustacean, but not molluscs.

Another crustacean allergen involved in invertebrate muscle contraction is SCBP (20 – 24 kDa), through binding of calcium ions [51, 52]. We identified two different SCBP contigs for each shrimp species (except endeavour shrimp), with PI values between 81-85%, implicating the presence of SCBP isoallergens. However, the significantly low abundance of SCBP_Contig_2 diminishes its role as an allergen compared to SCBP_Contig_1.

Other muscle regulatory protein identified include troponin. This protein is composed of three subunits, suffixed C, I, and T, with Troponin C and I being registered as allergens. TNC has been identified as an allergen in various crustaceans [53–55], cockroaches [56] and the storage mite [57]. Meanwhile, TNI has only been identified in narrow-clawed crayfish [58]. TIM, an enzyme that is involved in glucose metabolism, is also a registered allergen in north-sea shrimp [53], red swamp crayfish [59], american HDM [60], octopus [61] and wheat [62]. TNC, TNI, and TIM are highly conserved among shrimp species with sequence homology higher than 80%, 78%, and 87%, respectively.

Having considered the different shrimp allergens, it is important to note that apart from allergen presence, the abundance of isoforms also needs to be taken into consideration. Correlation between protein abundance and RNA-Seq data has been established, with some post-transcriptional cellular processes affecting this interpretation [63]. A study on European HDM allergen transcript levels using RNA-Seq data concluded that allergens have a higher level of abundance than non-allergens [64]; and their results were found to be to relatively similar to homologues identified in american HDM from a different study [65]. In particular, these dust mite allergen studies indicate that there is substantial correlation between RNA-Seq dependent abundance levels and a protein’s allergen status. Therefore, when there were more than one contig identified for any allergens in the five shrimps, we proceeded to analyse the isoform that had the highest abundance. Comparing known allergens within every shrimp species, SCBP, AK, and MLC were the most abundant in all five shrimps. Interestingly, TM was found to be significantly less abundant than SCBP (in king and endeavour shrimp); AK (in black tiger shrimp); and all three allergens (in whiteleg shrimp). However, TM is the major and most recognised shrimp allergen, despite its relatively low abundance. The stronger allergenicity is possible due to being very heat stable and having linear IgE binding epitopes as compared to AK, SCBP, or MLC [8, 28, 33].

In addition to previously implicated crustacean allergens, this study also identified up to 38 previously unreported but likely allergens (>50% PI), including seven proteins which are very-likely allergens (>70% PI). Three of these proteins, HSP70, alpha-tubulin, and chymotrypsin, have very high matches to known mite allergens in this study. Additionally, these three proteins have been identified as allergens in different mite and insect species [25, 66–69]. Subsequently, clinical cross-reactivity has been reported as crustacean-mite-insect syndrome [20, 38, 40, 42, 70–72], and we report here the most likely allergens involved. Furthermore, this study identified for the first time very likely allergens, responsible for possible cross-reactivity between shrimp and fish. Beta-enolase and aldolase A, enzymatic proteins that play a role in the glycolytic pathway [15], have previously been identified as heat labile allergens in various fish species and also chicken [73, 74]. Our findings implicate the importance of both proteins as strong candidate allergens in shrimps. In addition, other proteins that are identified to be candidate allergens include cyclophilin and G3PD. Cyclophilin allergen is generally found in fungi, plants and dust mites, and has been shown to have high rates of IgE-binding [60, 75]. Meanwhile, G3PD, an enzymatic protein that is involved in the process of glycolysis similar to aldolase A and beta-enolase, has been identified as allergens in wheat [76], and recently in cockroach and fish [25].

In conclusion, this study accomplished the comparative analyses of all known shrimp allergens derived from five different shrimp species’ transcriptomes assembled *de novo* from raw RNA-Seq data. The identification of previously known shrimp allergens validated the comprehensive approach utilised in this study. Moreover, over 30 additional proteins known for their allergenic properties in mite, fungi, plants, insect and fish were identified as candidate shrimp allergens. These includes HSP70, alpha-tubulin, chymotrypsin, beta-enolase aldolase A, cyclophilin and G3PD, which were further identified as very-likely candidate allergens of shrimps. Further immunological studies would be required to confirm clinical allergenicity in patients. The findings of this study will enable improved diagnostics for shrimp allergy and future therapeutics for this lifelong disease.

## Materials and methods

### Sample selection

Specimen of the five species of shrimps (*Litopenaeus vannamei, Penaeus monodon Fenneropenaeus merguiensis, Melicertus latisulcatus*, and *Metapenaeus endeavouri*) were supplied by the Commonwealth Scientific and Industrial Research Organisation (CSIRO) based in Queensland, Australia. *L. vannamei* and *P. monodon* samples originated from aquaculture farms whilst the other three species were caught as part of the CSIRO Northern Prawn Fishery Surveys from the benthic trawls in the Gulf of Carpenteria, Australia [77]. The shrimps were immersed in an ice-seawater slurry for a few minutes immediately after being caught, to be euthanized. Species-specific reference material were utilised to identify the species of shrimps [78]. Muscle tissue was then removed and stored in RNAlater^™^ (Invitrogen, Carlsbad, CA, USA) [79]. *P. monodon* samples were collected as described by Huerlimann et al (2018) [80]. Total RNA was extracted from the muscle tissue of three randomly selected adult shrimps of each of the five shrimp species (total of 15 samples) with an RNeasy Universal Extraction kit (QIAGEN) using manufacturer’s instruction in an RNase-free laboratory [79]. RNA concentration, quality and purity was assessed using a Nanodrop UV spectrophotometer (Thermo Fisher Scientific) and Agilent Bioanalyzer (Agilent Technologies), before being selected for sequencing.

### Illumina library preparation and RNA sequencing

All 15 samples were sequenced via Illumina HiSeq® 2500 System (Illumina Australia and New Zealand, VIC, Australia). Before sequencing, samples were quality checked with the Bioanalyzer RNA 6000 nano reagent kit (Agilent); and Illumina libraries were prepared using the TruSeq Stranded mRNA Library Preparation Kit (Illumina) according to established protocols at the Australian Genome Research Facility (AGRF). The resulting libraries were checked again with the TapeStation DNA 1000 TapeScreen Assay (Agilent). Cluster generation was performed immediately before sequencing on a cBot with HiSeq® PE Cluster Kit v4 – cBot. The sequencing was conducted using a HiSeq® SBS Kit on a HiSeq® 2500, operating with HiSeq Control Software v2.2.68 and base-calling with RTA v1.18.66.3. Raw RNA-Seq short read data for all samples are freely available on NCBI under BioProject PRJNA482687.

### De novo transcriptome assembly and quality control

RNA-Seq reads for all 15 samples were corrected using the software Rcorrector (v1.0.2) [81]. Transcriptomes of all 15 samples were individually assembled from their RNA-Seq data, *de novo*. The assembly was carried out using Trinity (v2.4.0) [82, 83]. The quality of the *de novo* transcriptome assembly was assessed using TransRate (v1.0.3) [84] and BUSCO (Benchmarking Universal Single-Copy Orthologs) (v1.2) [85] using the arthropoda odb9 database [86]. The quality score, also known as the TransRate score, is a score between 0.0 – 1.0 that is obtained by multiplying the mean of individual contig scores by the proportion of read pairs (original sequencing reads) that supported the transcriptome [84, 87]. The results of BUSCO assessment are given in percentages of complete (C), fragmented (F) and missing (M) genes within the transcriptome [85]. Using *L. vannamei* as an example, stepwise methods of sample extraction, sequencing, *de novo* transcriptome assembly and quality check are summarised and schematically represented in Figure 1.A.

### Removal of inconclusive dataset

Using the Rcorrected reads in an Assembly and Alignment-Free (AAF) method to create a phylogeny [88], it was discovered that one replicate of *M. latisulcatus* grouped with *M. endeavouri* rather than with the other two replicates of *M. latisulcatus*. To confirm the potentially misidentified sample, the assembled transcriptome was BLAST searched against the other *M. latisulcatus* and *M. endeavouri* transcriptomes, where the potentially misidentified sample also showed more similarity to *M. endeavouri*. Lastly, the transcriptomes were compared to known sequences of Enolase [89], which also confirmed that the misidentified sample is not *M. latisulcatus*.

### Allergen reference database construction

Known allergen AA sequences were retrieved from two reputable and peer-reviewed online databases to construct a reference allergen database for this study. The first is the World Health Organization & International Union of Immunological Societies (WHO/IUIS) Allergen Nomenclature database (www.allergen.org) [25]. The second is the AllergenOnline: Home of the FARRP (Food Allergy Research and Resource Program) Allergen Protein database (v.17) (www.allergenonline.org) [26, 27]. At the time of retrieval, the WHO/IUIS Allergen Nomenclature database contained 875 allergen AA sequences while the AllergenOnline database contained 2,035 allergen AA sequences [25–27]. After removing duplicates between the 2 databases, a total of 2,172 allergen AA sequences were compiled to form the reference allergen database for this study.

### BLAST search for allergens

The allergen database and the assembled transcripts for all 15 samples were imported into the Geneious™ software (v8.1.9, Biomatters Limited, USA) [90]. In order to compare and search for transcripts which contain similar sequences to the allergen sequences compiled in the allergen database, blastx searches were carried out using the BLAST (Basic Local Alignment Search Tool) module within the Geneious™ software. The criteria for the search conducted are shown in Supplementary Table 1.

### Refining the BLAST search results

The BLAST search results were filtered for matched sequences with a PI of 50% or more. Subject coverage (percentage of the allergen sequence that is covered by the matching transcript from the transcriptome) was manually calculated using the formula: Subject coverage = Sequence length / Subject length x 100%, where Sequence length is the length of the matched consensus sequence and the Subject length is the actual length of the allergen sequence from the constructed database. Results were then filtered again by selecting only sequences that have 90% or more subject coverage.

Duplicates of allergen sequences that aligned with contigs within the transcriptome were removed by keeping the top-matched allergen-transcript consensus sequence. The BLAST search results of 3 replicates of each species were then combined to form one list of allergens for every species and the duplicates (between replicates) were removed. Stepwise methods of allergen database construction and the processing of transcriptome data such as BLAST search, results refinement and removal of duplicates are schematically represented in Figure 1, using the three assembled transcriptome replicates of *L. vannamei* as an example.

### Analysing the BLAST search results

For each shrimp species, the matched allergen AA sequences were grouped into: ‘Shellfish’, ‘Mites’, ‘Insects’, ‘Fungi’, ‘Plants’, ‘Fish’, and ‘Other’, based on the organism that the allergen was documented in. The proportion of allergen sequences belonging to each group were graphed into a pie chart using GraphPad Prism version 7.03 for Windows [91] to show their distribution amongst different groups of allergen sources.

Multiple sequence alignment was conducted on all the contigs/transcripts that matched tropomyosin allergen in all five transcriptomes with shellfish tropomyosin allergens’ sequences (as reference). Mites’ and cockroaches’ tropomyosin allergen sequences were also included in the multiple sequence alignment that was conducted in Jalview2.1 using Clustal Omega [92]. Comparative AA sequence identities were carried out between the contigs from all five shrimp species that matched with tropomyosin, and previously reported crustacean, mites, and cockroach tropomyosin allergens using Clustal Omega, EMBL-EBI [93]. The multiple sequence alignment and comparative sequence identities were carried out for other documented crustacean allergens: arginine kinase, myosin light chain, sarcoplasmic calcium-binding protein, troponin C, troponin I, and triosephosphate isomerase.

Non-crustacean allergens that have a PI value of more than 70% were shortlisted as highly likely candidates of unreported allergens in shrimp species. These unreported allergens were selected based on their match with the transcriptome of a minimum of 70% PI in at least one of the 5 shrimp species.

### Measuring the abundance of allergen sequences

Abundance of each transcript/contigs within the transcriptomes, in transcript-per-million (TPM) values, was quantified using Salmon software [94]. Briefly, Salmon is a software that estimates the abundance of each contig by measuring the number of reads from the RNA-Seq data that align to the contig being measured [94]. Abundance estimation values for all known crustacean allergens were retrieved from all 15 samples. For each allergen in each sample, the estimated abundance value is the sum of all TPM values of all the contigs that matched with that allergen. The mean TPM values with standard deviation error bars for each allergen of the three replicates for each shrimp species are graphically represented in Figure 7 and 8. Standard deviation error bars were omitted from *M. latisulcatus* samples as only 2 replicates were investigated in this study. We first analysed the difference in abundance of all contigs representing a specific allergen, between the 5 shrimp species (Figure 7). In order to look for significant differences between two contigs representing the same allergen, we used unpaired *T-test* using GraphPad Prism version 7.03 for Windows [91]. Next, we analysed the difference in abundance of allergens within each shrimp species (Figure 8). For this analyses, we only took into account the contig with the highest abundance, when there are more than one contig representing one allergen. To analyse significant differences between the seven crustacean allergens’ abundance, we used One-way ANOVA test using GraphPad Prism version 7.03 for Windows [91].

### Molecular phylogenetic tree building of TM, AK, MLC and SCBP

Published AA sequences of the four widely studied crustacean allergens, TM, AK, MLC and SCBP belonging to edible crustacean and mollusc species; and allergy causing mite and insect species were mined from NCBI Genbank and UniProt databases. The proteins which are not registered as an allergen in WHO/IUIS or AllergenOnline databases were also included. To determine the evolutionary distance between the same proteins from different species, molecular phylogenetic trees for each protein was built using MEGA X software (v10.0.5). The trees were constructed using the neighbour-joining method with the Poisson correction model. Hence, the branch lengths are the proportion of AA substitutions per site. Bootstrap test was also included (10,000 replicates) and the percentages are shown next to the branches. The gaps which occurred in alignment were treated as pairwise deletion.

## Supporting information

All Supplemental Figures and Tables

## Authors’ contributions

EJ and AL conceptualized the research objectives. SK, RH, EJ, RN, AT, and AL designed the research methods. RH, NW, and DJ provided the shrimp samples, conducted the RNA extraction and funded the RNA sequencing. SK and RH carried out the *de novo* transcriptome assembly, transcriptome quality check, BLAST search and relative abundance estimation. SK and RN constructed the allergen reference database and the molecular phylogenetic tree of known shellfish allergens. SK, RN, EJ, TR, AT, SDK, and AL determined the relevant allergens identified within the shrimp transcriptomes and the downstream analyses of these potential allergens. SK, RN, TR, RH, and AL designed the figures and tables. SK and AL wrote the first draft. All authors contributed to manuscript editing and revision.

## Supplementary material

**Supplementary Figure 1:** Comparison of amino acid sequence identities of (1-7) contigs from five shrimp species that matched with Troponin C (TNC) allergen, (8-9) known shrimp TNC allergen, and (10-14) cockroach and storage mite TNC allergen. The sequence identities were calculated using multiple sequence alignment in Clustal Omega (EMBL-EBI).

**Supplementary Figure 2:** Comparison of amino acid sequence identities of (1-5) contigs from five shrimp species that matched with Troponin I (TNI) allergen and (6) known crayfish TNI allergen. The sequence identities were calculated using multiple sequence alignment in Clustal Omega (EMBL-EBI).

**Supplementary Figure 3:** Comparison of amino acid sequence identities of (1-5) contigs from five shrimp species that matched with triosephosphate isomerase (TIM) allergen, (6) known shrimp TIM allergen, and (7-8) house dust mite TIM allergen. The sequence identities were calculated using multiple sequence alignment in Clustal Omega (EMBL-EBI).

**Supplementary Figure 4:** Multiple sequence alignment of (1-2) known shrimp tropomyosin (TM) allergen, (3-9) contigs from five shrimp species that matched with TM allergen and (10-12) TM allergen sequences from house dust mite and cockroaches. Multiple sequence alignment was conducted in Jalview 2.1 using Clustal Omega.

**Supplementary Figure 5:** Multiple sequence alignment of (1-2) known shrimp arginine kinase (AK) allergen, (3-8) contigs from five shrimp species that matched AK allergen and (9-12) AK allergen sequences from house dust mites and cockroaches. Multiple sequence alignment was conducted in Jalview 2.1 using Clustal Omega.

**Supplementary Figure 6:** Multiple sequence alignment of (1-3) known shrimp myosin light chain (MLC) allergen, (4-8) contigs from five shrimp species that matched with MLC allergen and (9-10) house dust mite and cockroach MLC allergen. Multiple sequence alignment was conducted in Jalview 2.1 using Clustal Omega.

**Supplementary Figure 7:** Multiple sequence alignment of (1-3) known shrimp sarcoplasmic calcium-binding protein (SCBP) allergen and (4-12) contigs from five shrimp species that matched with SCBP allergen. Multiple sequence alignment was conducted in Jalview 2.1 using Clustal Omega.

**Supplementary Figure 8:** Multiple sequence alignment of (1-2) known shrimp Troponin C (TNC) allergen, (3-9) contigs from five shrimp species that matched with TNC allergen and (10-14) TNC allergen sequences from house dust mites and cockroaches. Multiple sequence alignment was conducted in Jalview 2.1 using Clustal Omega.

**Supplementary Figure 9:** Multiple sequence alignment of (1) known crayfish Troponin I (TNI) allergen and (2-6) contigs from five shrimp species that matched with TNI allergen. Multiple sequence alignment was conducted in Jalview 2.1 using Clustal Omega.

**Supplementary Figure 10:** Multiple sequence alignment of (1) known shrimp Triosephosphate isomerase (TIM) allergen, (2-6) contigs from five shrimp species that matched with TIM allergen and (7-8) TIM allergen sequences from house dust mites. Multiple sequence alignment was conducted in Jalview 2.1 using Clustal Omega.

**Supplementary Table 1: Criteria used in the BLAST search conducted.** The criteria shown here are only for the BLAST search utility within the Geneious™ software. Additional search criteria (for this project) was later used in the refining process of the search results, e.g. Minimum % Pairwise Identity of 50%.

