## Supplementary material for "Novel allergen discovery through comprehensive *de novo* transcriptomic analyses of 5 shrimp species": All Supplemental Figures and Tables

**Title page**

**Journal:** Genomics, Proteomics, and Bioinformatics

**Running title:** *Karnaneedi et al / Transcriptomic analysis of shrimp allergens*

**Supplementary material**

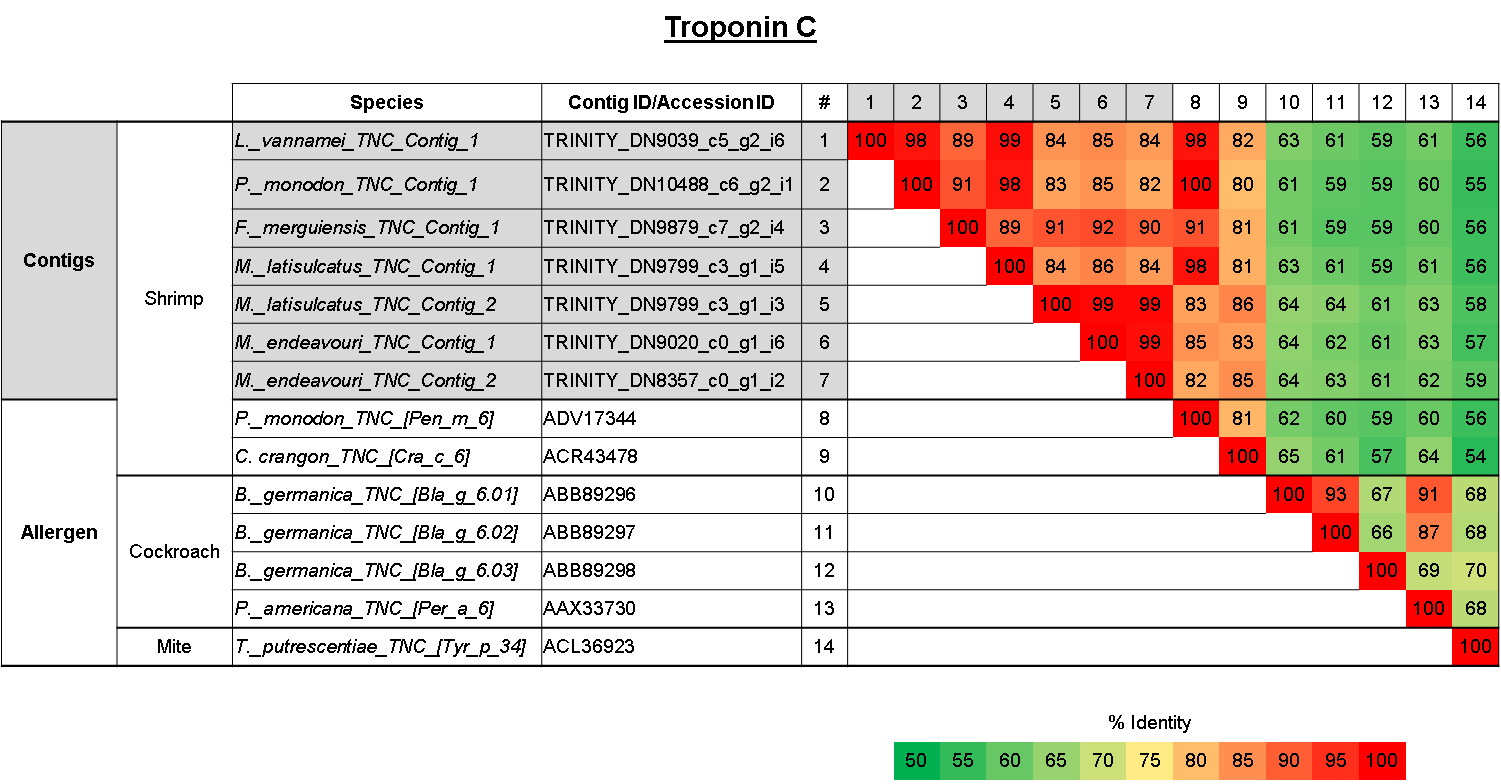

**Supplementary Figure 1:** Comparison of amino acid sequence identities of (1-7) contigs from five shrimp species that matched with Troponin C (TNC) allergen, (8-9) known shrimp TNC allergen, and (10-14) cockroach and storage mite TNC allergen. The sequence identities were calculated using multiple sequence alignment in Clustal Omega (EMBL-EBI).

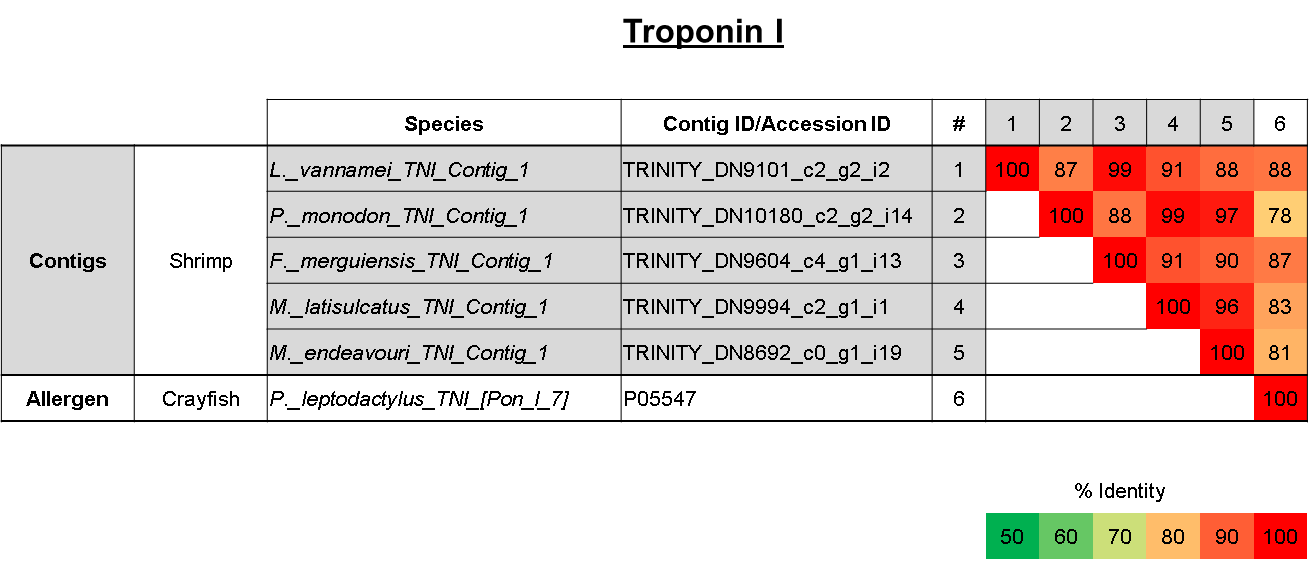

**Supplementary Figure 2:** Comparison of amino acid sequence identities of (1-5) contigs from five shrimp species that matched with Troponin I (TNI) allergen and (6) known crayfish TNI allergen. The sequence identities were calculated using multiple sequence alignment in Clustal Omega (EMBL-EBI).

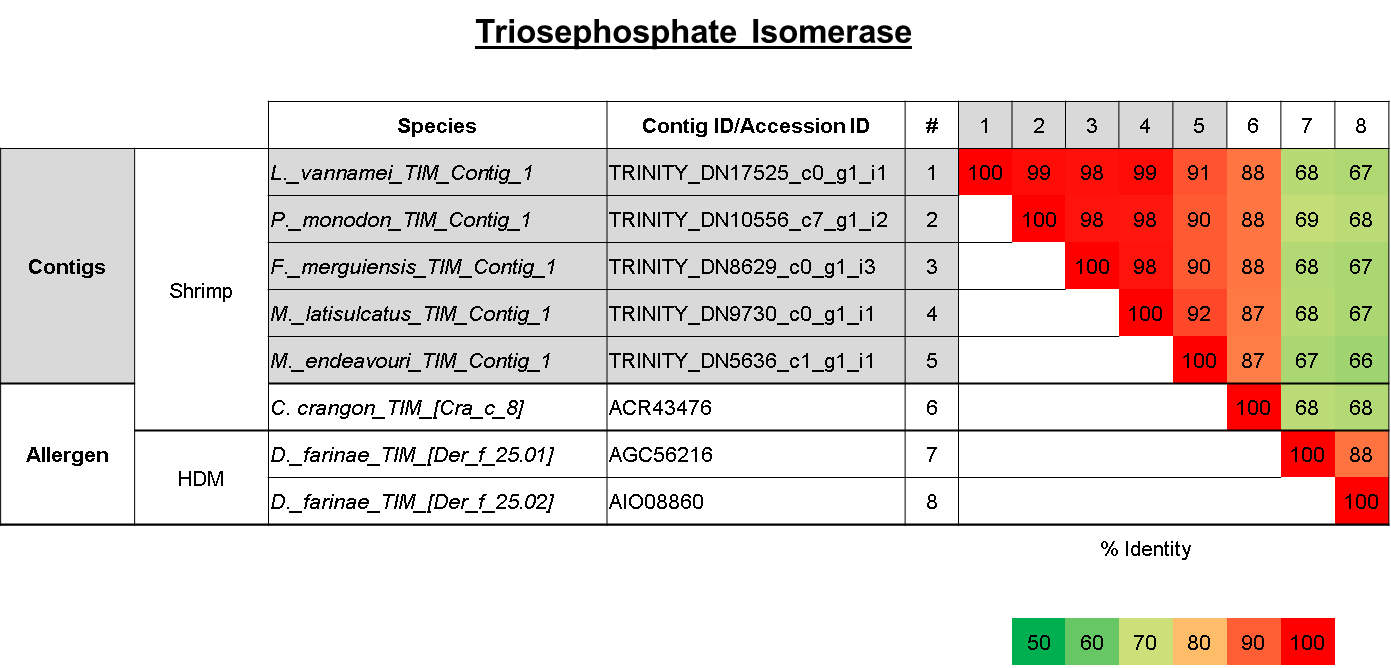

**Supplementary Figure 3:** Comparison of amino acid sequence identities of (1-5) contigs from five shrimp species that matched with triosephosphate isomerase (TIM) allergen, (6) known shrimp TIM allergen, and (7-8) house dust mite TIM allergen. The sequence identities were calculated using multiple sequence alignment in Clustal Omega (EMBL-EBI).

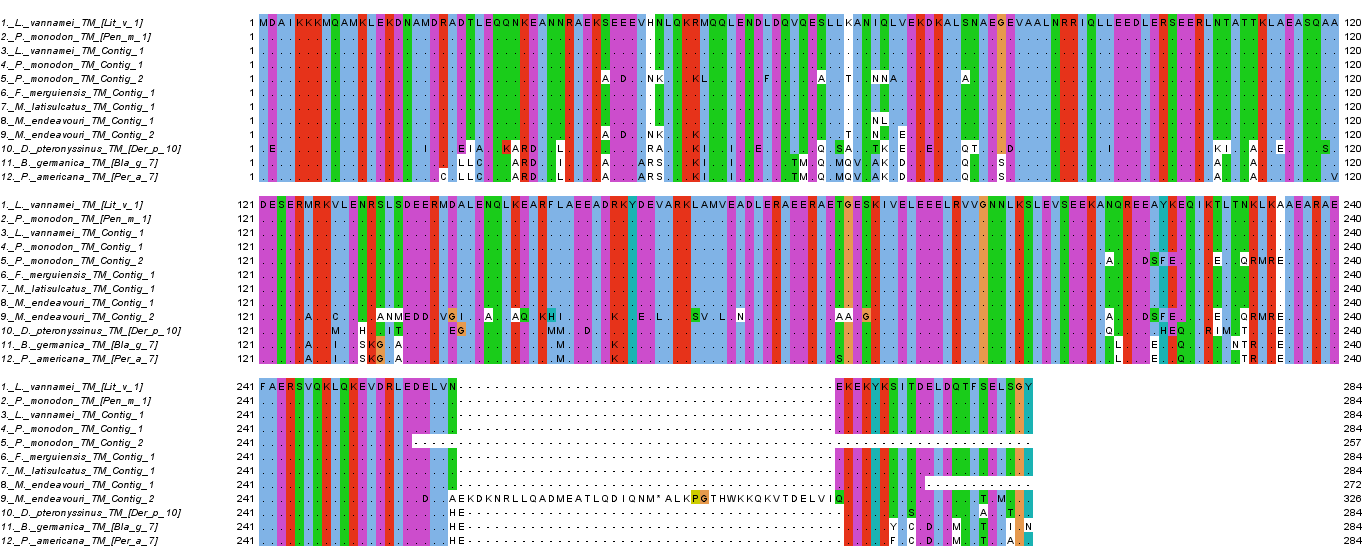

**Supplementary Figure 4:** Multiple sequence alignment of (1-2) known shrimp tropomyosin (TM) allergen, (3-9) contigs from five shrimp species that matched with TM allergen and (10-12) TM allergen sequences from house dust mite and cockroaches. Multiple sequence alignment was conducted in Jalview 2.1 using Clustal Omega.

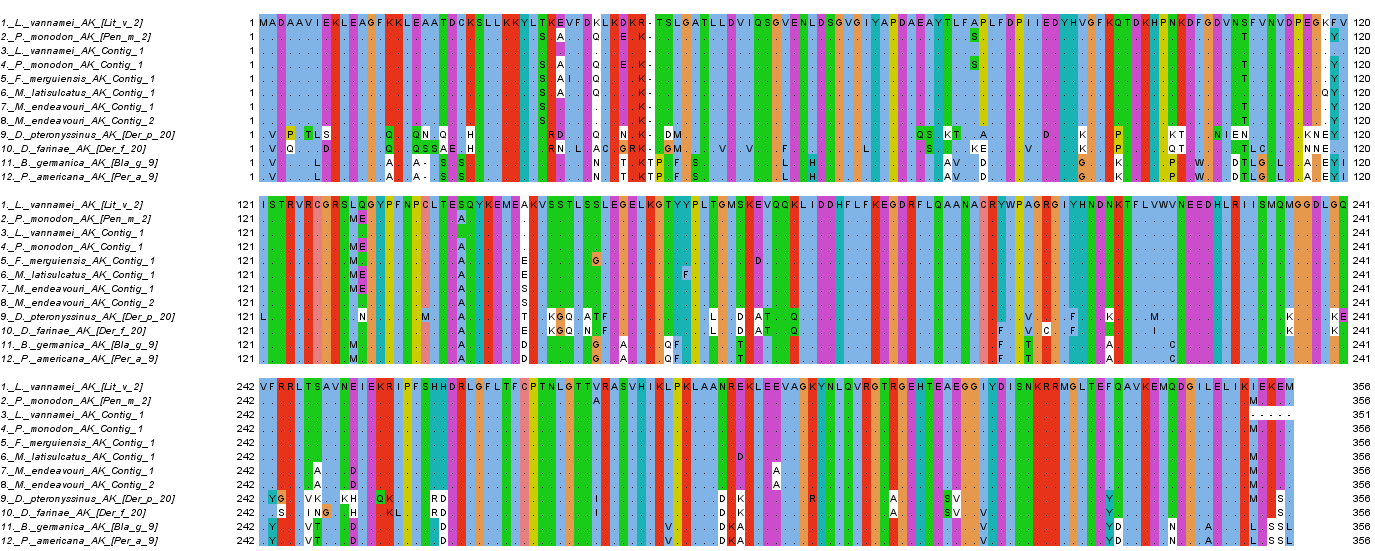

**Supplementary Figure 5:** Multiple sequence alignment of (1-2) known shrimp arginine kinase (AK) allergen, (3-8) contigs from five shrimp species that matched AK allergen and (9-12) AK allergen sequences from house dust mites and cockroaches. Multiple sequence alignment was conducted in Jalview 2.1 using Clustal Omega.

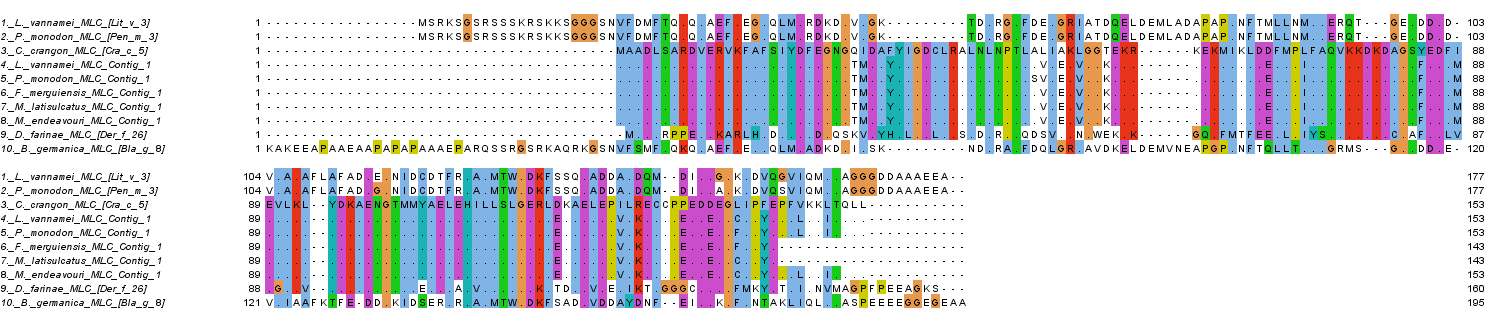

**Supplementary Figure 6:** Multiple sequence alignment of (1-3) known shrimp myosin light chain (MLC) allergen, (4-8) contigs from five shrimp species that matched with MLC allergen and (9-10) house dust mite and cockroach MLC allergen. Multiple sequence alignment was conducted in Jalview 2.1 using Clustal Omega.

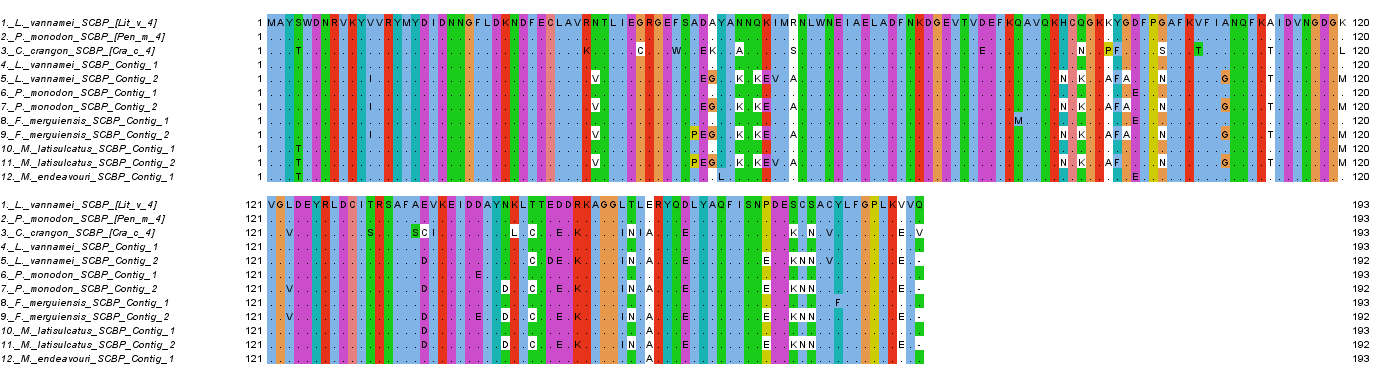

**Supplementary Figure 7:** Multiple sequence alignment of (1-3) known shrimp sarcoplasmic calcium-binding protein (SCBP) allergen and (4-12) contigs from five shrimp species that matched with SCBP allergen. Multiple sequence alignment was conducted in Jalview 2.1 using Clustal Omega.

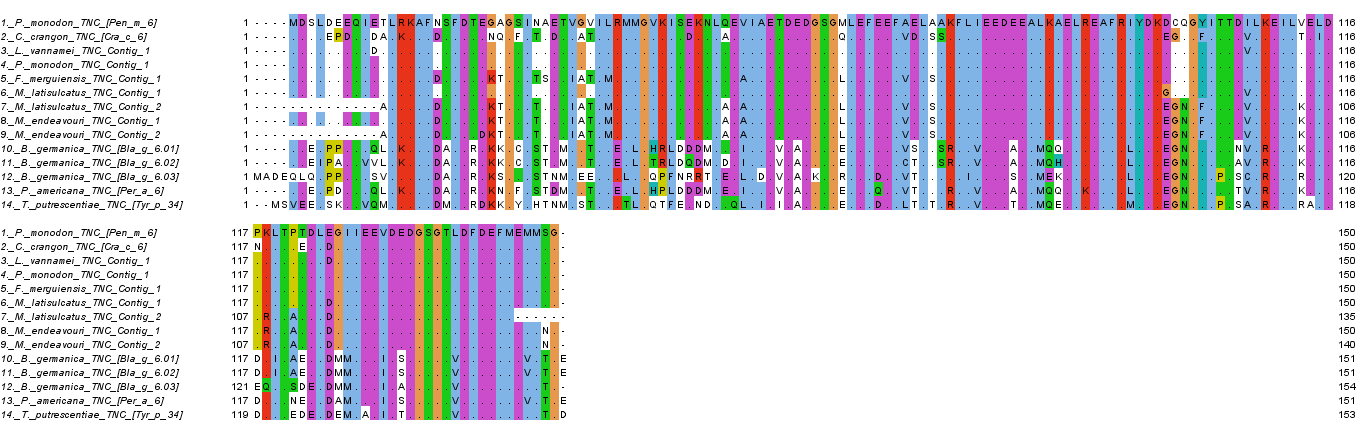

**Supplementary Figure 8:** Multiple sequence alignment of (1-2) known shrimp Troponin C (TNC) allergen, (3-9) contigs from five shrimp species that matched with TNC allergen and (10-14) TNC allergen sequences from house dust mites and cockroaches. Multiple sequence alignment was conducted in Jalview 2.1 using Clustal Omega.

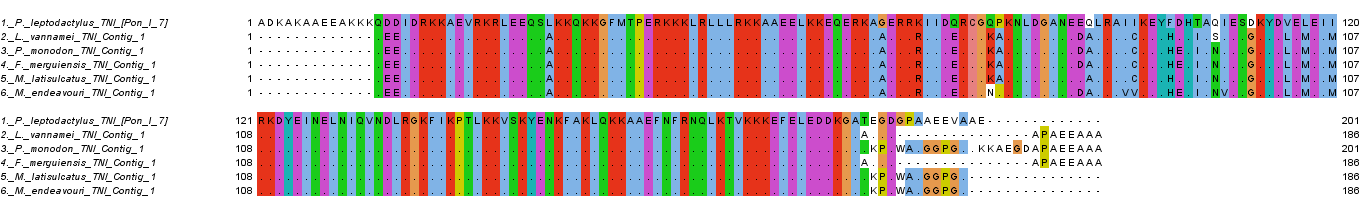

**Supplementary Figure 9:** Multiple sequence alignment of (1) known crayfish Troponin I (TNI) allergen and (2-6) contigs from five shrimp species that matched with TNI allergen. Multiple sequence alignment was conducted in Jalview 2.1 using Clustal Omega.

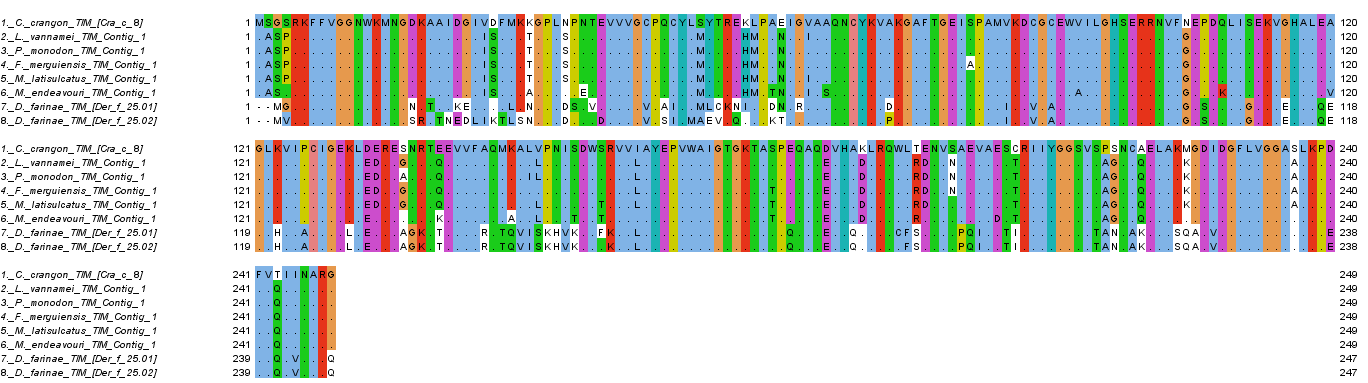

**Supplementary Figure 10:** Multiple sequence alignment of (1) known shrimp Triosephosphate isomerase (TIM) allergen, (2-6) contigs from five shrimp species that matched with TIM allergen and (7-8) TIM allergen sequences from house dust mites. Multiple sequence alignment was conducted in Jalview 2.1 using Clustal Omega.

**Supplementary Table 1:** **Criteria used in the BLAST search conducted.**
The criteria shown here are only for the BLAST search utility within the Geneious™ software. Additional search criteria (for this project) was later used in the refining process of the search results, e.g. Minimum % Pairwise Identity of 50%.

| BLAST search criteria | |
| --- | --- |
| Query | Batch search of nucleotide sequences |
| Database | Allergens (AA) |
| Program | blastx |
| Results | Hit table |
| Retrieve | Matching regions with annotations |
| Maximum Hits | 1 |
| Low complexity filter | “*checked*” |
| Max E-value | 1e-7 |
| Word Size | 3 |
| Matrix | BLOSUM62 |
| Number of CPUs | 30 |
| Gap cost (Open Extend) | 11 1 |
